# Fourier Ring Correlation Simplifies Image Restoration in Fluorescence Microscopy

**DOI:** 10.1101/535583

**Authors:** Sami Koho, Giorgio Tortarolo, Marco Castello, Takahiro Deguchi, Alberto Diaspro, Giuseppe Vicidomini

## Abstract

Fourier ring correlation (FRC) has recently gained some popularity among (super-resolution) fluorescence microscopists as a straightforward and objective method to measure the effective resolution of a microscopy image. While the knowledge of the numeric resolution value is helpful in e.g. interpreting imaging results, much more practical use can be made of FRC analysis – in this article we propose novel blind image restoration methods enabled by it. We apply FRC to perform image de-noising by frequency domain filtering. We propose novel blind linear and non-linear image deconvolution methods that use FRC to estimate the effective point-spread-function, directly from the images, with no need for prior knowledge of the instrument or sample characteristics. The deconvolution is shown to work exquisitely with both two- and three-dimensional images. We also show how FRC can be used as a powerful metric to observe the progress of iterative deconvolution. While developing the image restoration methods, we also addressed two important limitations in FRC that are of more general interest: how to make FRC work with single images and with three-dimensional images with anisotropic resolution.

---

Reliable and realistic estimation of spatial resolution in fluorescence microscopy images has for decades been a subject of passionate scientific debate. With the development of fluorescence nanoscopy techniques [1] this debate has resurfaced with new-found fervour, e.g. [2, 3].

Typically resolution is estimated by measuring either the minimum resolvable distance between two adjacent structures in an image – as per the classical Rayleigh/Abbe/Sparrow resolution definitions – or, alternatively it can be estimated from the full-width-half-maximum (FWHM) width of sub-resolution spatial structures. In order to perform either one of the two measurements, suitable structures need to be subjectively identified and manually measured; ideally, the measurements should be repeated at several positions to gain some statistical basis for the estimate. This task is both tedious as well as error-prone.

Fourier ring correlation (FRC) [4, 5] and Fourier shell correlation (FSC) [6] – essentially, FRC generalized to 3D – have for decades been used to estimate image resolution in electron cryomicroscopy (CryoEM). Recently FRC was adapted for optical nanoscopy, by us [7] and others [8, 9], to address the issues with the traditional resolution assessment methods. It is based on a normalized cross-correlation histogram measure calculated in the frequency domain between two images of the same region-of-interest, with independent noise realizations. For FRC/FSC calculation, the spatial frequency spectra of the two images are divided into bins, which produces a series of concentric rings/shells in the polar-form frequency domain images (hence the names). The FRC/FSC histogram is formed by calculating a correlation value for each bin according to

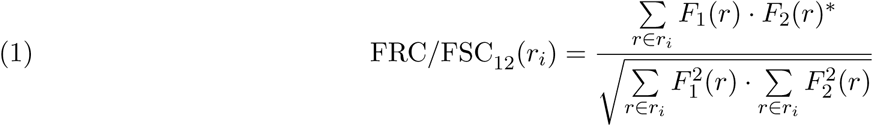

, where *F*_1_ and *F*_2_ are the Fourier transforms of the two images and *r_i_* the *i*th frequency bin. The image resolution in FRC/FSC is defined from the histogram, as a cut-off frequency at which the cross-correlation value drops below a pre-set threshold value. Advantages of the FRC/FSC are that it is fully automatic, quantitative and depends both on sample and microscope characteristics – it is also less prone to subjective bias and measurement errors, although the choice of the appropriate resolution threshold criterion still requires some input from the researcher, as no single solution seems to be correct in all applications [10, 8].

While the resolution estimation certainly is an interesting application in itself, in our view, the true potential of FRC/FSC is in much more practical tasks. In this paper we show several examples of advanced image restoration methods that leverage FRC/FSC measures. We apply FRC to perform image de-noising by frequency domain filtering. We propose novel blind linear and iterative image deconvolution methods that use FRC/FSC measurements to estimate the effective point-spread-function (PSF) of the microscope, directly from the images, with no need for prior knowledge of the instrument characteristics. The deconvolution is shown to work exquisitely with both two- and three-dimensional (2D/3D) images. We also show how FRC can be used as a powerful metric to observe the progress of iterative deconvolution tasks.

## Results

### One-image FRC/FSC measures

For traditional FRC/FSC analysis two images of the very same region-of-interest and with independent noise realisations are needed. These two images can be obtained in a point-scanning microscope by using sequential imaging on a line-by-line base. Compared to frame-by-frame sequential imaging, line-by-line allows overcoming potential differences between the two images induced by focus drift, sample dynamics, or photobleaching. Similarly, sequential imaging on a pixel-by-pixel, pulse-by-pulse and photons-by-photons basis can further decrease the differences between the two images, but needs more sophisticated (e.g. time-resolved) instrumentation [7]. On the other hand, frame-by-frame sequential imaging is the only option for wide-field microscopy, but high frame-rate is needed to ensure the sameness of the two images, particularly with live samples. Especially with the image processing tasks in mind, in which the requirement of two images introduces a unnecessary computational overhead, and to make the FRC/FSC analysis compatible with any microscopy architecture, we developed a method to calculate FRC/FSC from a single image (one-image FRC/FSC).

In order to perform FRC/FSC analysis on a single image, one needs to find a way to form statistically independent image subsets that share the same details, but different noise realizations, by some form of sub-sampling. As described in (Figure 1a)) we propose to do this by dividing a single image into four subsets, i.e., two image pairs. The first pair is formed by taking every pixel with (even, even) row/column indexes to form one sub-image and (odd, odd) indexes to the other. The second image pair is formed from pixels at (even, odd) and (odd, even) indexes. FRC can be calculated from either one of the image pairs alone, but we noticed that averaging two measurements helps to deal with special spectral domain symmetries (Figure S. 1). With 3D images (FSC) the same splitting method is used, except that in the axial direction (z) layers are summed pairwise to maintain image proportions; we get back to the 3D measurements later.

The dimensions of the four sub-images are identical, exactly half of the size of the original image. Because sub-sampling inevitably leads to loss of information, one might be keen to think that such a method is only feasible on significantly oversampled data, i.e. with sampling much higher than Nyquist, 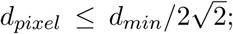; *d_pixel_* for pixel size; *d_min_* for expected image resolution; we use the 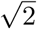 factor in our Nyquist limit definition to ensure sufficient sampling in all directions, assuming square pixels on a rectangular sampling grid [11]. However, as illustrated by the diagonal lines in (Figure 1a)), the proposed sub-sampling pattern introduces an offset of 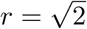 pixels between the two sub-images in each of the two sets, which as described in detail in (Note S. 1), demonstrates itself as a rather interesting modulation in the frequency domain that has the property of compressing the FRC/FSC curve – by compression one means that details are shifted to a lower frequency than where they actually should be. We simply compensate for the shift by re-scaling the frequency axis of the FRC/FSC curves by the 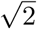 factor, which makes it possible to use the single image splitting method with images much more sparsely sampled than predicted above, in practice up-to the Nyquist limit.

In order to properly understand how the single image FRC/FSC works, we ran a series of test measurements. In each of the tests, the regular two-image FRC with 1/7 threshold, as proposed in [8, 7], was used as reference. In one-image FRC the resolution threshold corresponding to SNR_*e*_=0.25 [10] was used. In (Figure 1b)) we compare one- and two-image FRC results from a dataset consisting of a series of confocal fluorescence microscopy images of a vimentin stained cell, acquired in a single field-of-view, with different pixel sizes. Continuous autofocus was enabled on the microscope to minimize axial drift. The effective resolution increases slightly towards smaller pixel sizes, because the pixel dwell time was kept fixed at 10*µs* in all measures. From the results it is evident that the one-image and two-image FRC curves perform rather differently – the one-image curve decays more slowly. For this reason, with extremely small pixel sizes (29nm, *d_min_/*8) the one-image method with the SNR_*e*_=0.25 threshold appears to somewhat overestimate the resolution (*circa* 25nm difference). On the other hand at pixel sizes approaching the Nyquist limit (i.e. 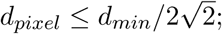; assuming a square sampling grid), the SNR_*e*_=0.25 threshold seems to produce somewhat conservative resolution values – in fact the 1/7 would seem more appropriate. It is important to note that the only pixel size at which the one-image FRC clearly does not work is 113nm, which is natural, as such a sparse sampling is below the above Nyquist limit, and causes undersampling and possible aliasing in the diagonal direction. To further confirm the appropriateness of the 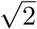 factor frequency axis correction, we simulated the same situation in the two-image FRC with the vimentin cell images, by applying the offset (x=1 px, y=1 px) to one of the images in each image pair before FRC calculation (Figure S. 2). Interestingly nearly identical results with the ones shown in (Figure 1b)) with one-image measures were obtained. We then characterised the one-image FRC by analysing a series of stimulated-emission-depletion (STED)[12] super-resolution microscope images of nuclear pore complexes (Figure 1c)) with same pixel size but different resolution. The one- and two-image FRC results correlate well from confocal to near the Nyquist limit; once again the SNR_*e*_=0.25 threshold seems to be a bit conservative at the higher frequencies, but in general performs well. The high frequency behaviour of the one-image FRC was further analysed with two sets of STED images of 40nm Crimson fluorescent beads, with different pixel sizes (Figure S. 3): at the highest frequencies the one-image measure starts to fall off, but at worst produces a *circa* 10nm error. As a last step we wanted to confirm that the normalization of the FRC curve works as expected. In (Figure 1d)) FRC results from a dataset consisting of a series of confocal fluorescence microscopy images of a vimentin stained cell, acquired in a single field-of-view, with different illumination intensity levels are shown: both one- and two-image FRC are independent of the illumination intensity (as long as there is sufficient SNR to avoid resolution loss). To conclude, the one-image FRC was shown to work well with a wide range of sampling densities (*d_pixel_ ∈* [*d_min_/*3*, d_min_/*8]). With significantly over-sampled data, the SNR_*e*_=0.25 threshold appears to be a bit too low, but the errors remain minor. With such data, in order to match the numerical values with two-image FRC, a threshold (e.g. *≈* 0.4, when *d_pixel_ ≤ d_min_/*8) could be used. Notably, the range of sampling density valid for the one-image FRC/FSC analysis matches with the typical sampling for both wide-field and point-scanning microscopy. Furthermore, for *d_pixel_ ≥ d_min_/*3 the FRC/FSC analysis fails also in the two-image case, whilst for *d_pixel_ ≤ d_min_/*3 the image can be binned to satisfy the sampling condition.

**Figure 1:**
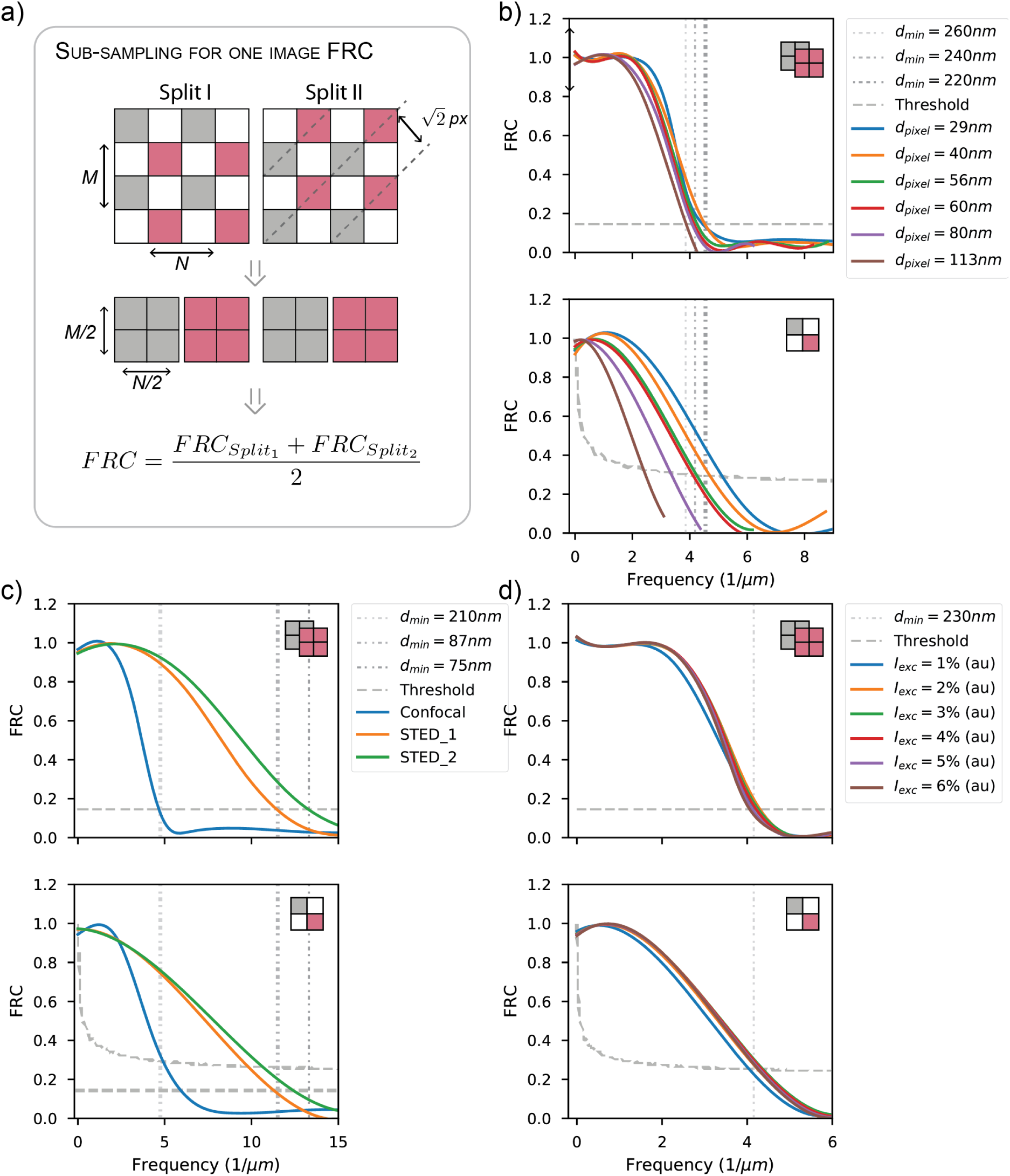
FRC measurements using a single image. In a) the principle of the single image sub-sampling for FRC is illustrated. Diagonally adjacent pixels are selected to form two sets of sub-images. Such splitting introduces a (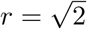) offset to the sub-images. FRC measures from the two-image sets are averaged to compensate for possible spectral domain asymmetries. In b) one- and two-image FRC results with a confocal dataset consisting of a pixel size gradient are shown. In c) one- and two-image FRC results with a STED microscope dataset with resolution gradient are shown. In d) one- and two-image FRC results with a confocal dataset with excitation intensity gradient are shown.

There are also a number of other ways that one might consider sub-sampling an image. In (Figure S. 4) all the alternatives that we came up with are shown. In (Figure S. 5) each method is compared with three different images. It is immediately evident that only the diagonal splitting works properly on all of the images. The other methods do not benefit from the bandwidth gain and also suffer from spatial biases and correlations produced by the asymmetric sampling patterns and summing. The various pros and cons are further summarized in (Note S. 4). As a further note, the benefit of averaging the two diagonal measures is highlighted in (Figure S. 5 c)), as the microtubulin stained HeLa cell image has a peculiar frequency domain symmetry (Figure S. 1), which produces a higher resolution value in one direction than the other - one could of course instead of averaging keep the higher of the two values. In absence of such special distribution of image details (Figure S. 5 a-b)) the results from the two diagonals are identical.

### Frequency domain filtering based on FRC cut-off

In (Figure 2) results obtained with three types of Fourier domain low-pass filters are shown, with a low-quality confocal image (low signal, strong background) of HeLa cell tubulin cytoskeleton (pixel size 51nm). The FRC analysis (one-dual and two-image) estimated a resolution of 0.2*µm*. The inverse of the resolution value was used as a threshold for the Ideal, Butterworth and Gaussian low-pass frequency domain filters. All the filters were able to significantly reduce the noise, with no apparent effect on the fine image details. The Ideal filter works excellently: it is able to practically remove all the high-frequency noise, with a faint low-frequency noise pattern visible in the background (from scanning, laser fluctuation etc.). The use of an Ideal filter is not typically recommended, as introducing such sharp edges into the frequency domain may produce artefacts (ringing effects) in the filtering results. No such effects can be seen in (Figure 2), supposedly because there are no visible details – and signal power is very close to zero, at frequencies higher than the FRC threshold. It should thus be rather safe to use an Ideal filter in this case. The two more gentle Frequency domain filters, Gaussian and Butterworth, also clearly work, but are not able to remove all the noise.

**Figure 2:**
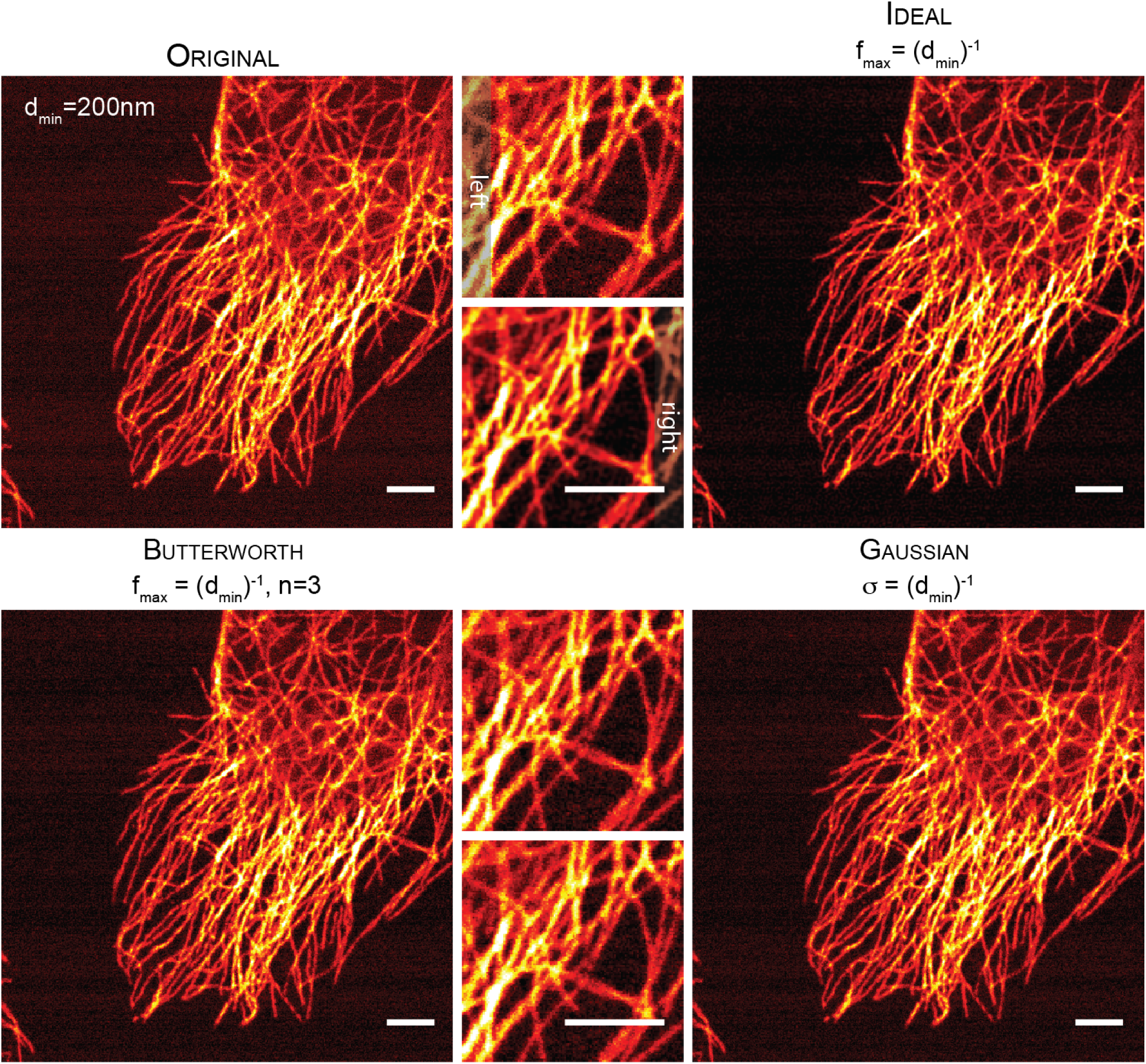
FRC-based Fourier domain low-pass filtering. FRC measurement based Fourier domain low-pass filtering approach is demonstrated on a noisy microtubulin stained HeLa cell image. The resolution value obtained with FRC measurement (*d_min_* = 200*nm*) is used as a cut-off frequency for the filters. An Ideal filter is shown to nearly completely remove high-frequency noise, whereas both Butterworth and Gaussian filters achieve less dramatic results. Scale bars 3*µm*

### FRC enabled blind Deconvolution

With deconvolution one tries to enhance the contrast and effective resolution in an image, by using information of the microscope’s transfer function – the PSF – as a prior knowledge in a image restoration algorithm. Here we leverage the one-image FRC, to estimate the PSF directly from images, with no need for prior knowledge of the instrument or sample characteristics. The PSF is based on a simple Gaussian model, in which the one-image FRC resolution (Figure 1) is used as FWHM value (FWHM = 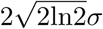).

In (Figure 3a)) results of 2D blind Wiener deconvolution (filtering) with PSF estimated with one-image FRC are shown. The same noisy microtubulin stained HeLa cell image that was used in the FFT filtering section is used in here. The Wiener deconvolution result shows dramatic enhancement of contrast and effective resolution – the latter is confirmed by FRC measurement of the deconvolution result. The regularization SNR value was selected subjectively to produce a sharp result with good contrast.

We then used the same FRC based PSF in an iterative Richardson-Lucy (RL) devonvolution algorithm. In our RL implementation, the FRC is calculated after each iteration with the method described in (Figure 1 a); in the *Adjustive RL* we also update the PSF after each iteration, but otherwise these measures are only used to assess the effective resolution of the deconvolution results, similar to what was done in in [13]. The blind RL deconvolution results with FRC based PSF (Figure 3b) show dramatically improved contrast and effective resolution. The FRC measures of various intermediate deconvolution estimates reveals that no major gains in term of resolution are made after approx. 10 iterations. The *≈* 170*nm* resolution that is reached at the end of the iteration is very near the confocal cut-off and certainly better than widefield (FWHM_*confocal*_ = 0.37 *⋅* 525*nm/*1.4 = 140*nm*; FWHM_*wf*_ = 0.51 ⋅ 525*nm/*1.4 = 190*nm*) [14], especially considering that the the one-image FRC at SNR_*e*_ = 0.25 threshold tends to produce conservative values near the Nyquist limit.

Usually in classical iterative blind deconvolution algorithms [15, 16] one starts with some sort of a rough PSF estimate that is updated at every iteration. We did the same in (Figure 3c) by updating the PSF after each iteration, based on one-image FRC measurement. While the *Adjustive RL* deconvolution appears to be completely stable, the FRC measures of the results show that compared to a) the iteration takes a longer time to converge, and the result after 40 iterations is also somewhat more noisy.

As a last step, we introduced a total variation (TV) regularization (*λ_T V_* = 5 *⋅* 10^*−*4^) [17] to the *Adjustive RL* algorithm, as it should help to control the slight overfitting that seemed to take place. The smoothing constraint does greatly enhance the contrast, but produces somewhat blurry looking results. The FRC measurements in (Figure 3a-d)) suggest that despite apparent differences, all the four algorithms produce results with nearly identical effective resolution; this is supported by line profile measurements in (Figure S. 8).

From the FRC measures shown in (Figure 3 b-d)), it is evident that the effective resolution in the deconvolution results tends to converge to a nearly fixed value after a number of iterations. This insight motivated us to look into another long-standing problem with RL and other iterative deconvolution algorithms: that one does not really know when the algorithm should be stopped. Quite commonly this is decided by trial and error, by running the algorithm with different iteration counts, and picking the best looking result. Several parameters have been proposed as well to observe the progress of deconvolution, but sadly, they are not very reliable or commonly used. In (Figure S. 9)b)) we show a plot for the effective resolution in (Figure 3 b)), as a function of deconvolution iterations. The blind RL result in (Figure S. 9)a)) is the same that was shown in (Figure 3b)). The effective resolution value reaches its maximum after 38 iterations, after then the resolution starts to slowly decrease. This is in good agreement with theory [18], since the exponential increase of noise, as a function of iteration count, eventually reduces the amount of signal and the effective resolution [19]. It is also evident from the *resolution* = *f*(*iteration*) curve that only very minor gains in resolution are made after *≈* 10 iterations. We therefore set up two stopping conditions for the deconvolution based on the first derivative of the *resolution* = *f*(*iteration*) curve: (i) the iteration of the first zero crossing, to indicate the maximum resolution, or (ii) the iteration after which the derivative decreases below a threshold (1*nm/iteration* shown as a horizontal line in (Figure S. 9)b)).

**Figure 3:**
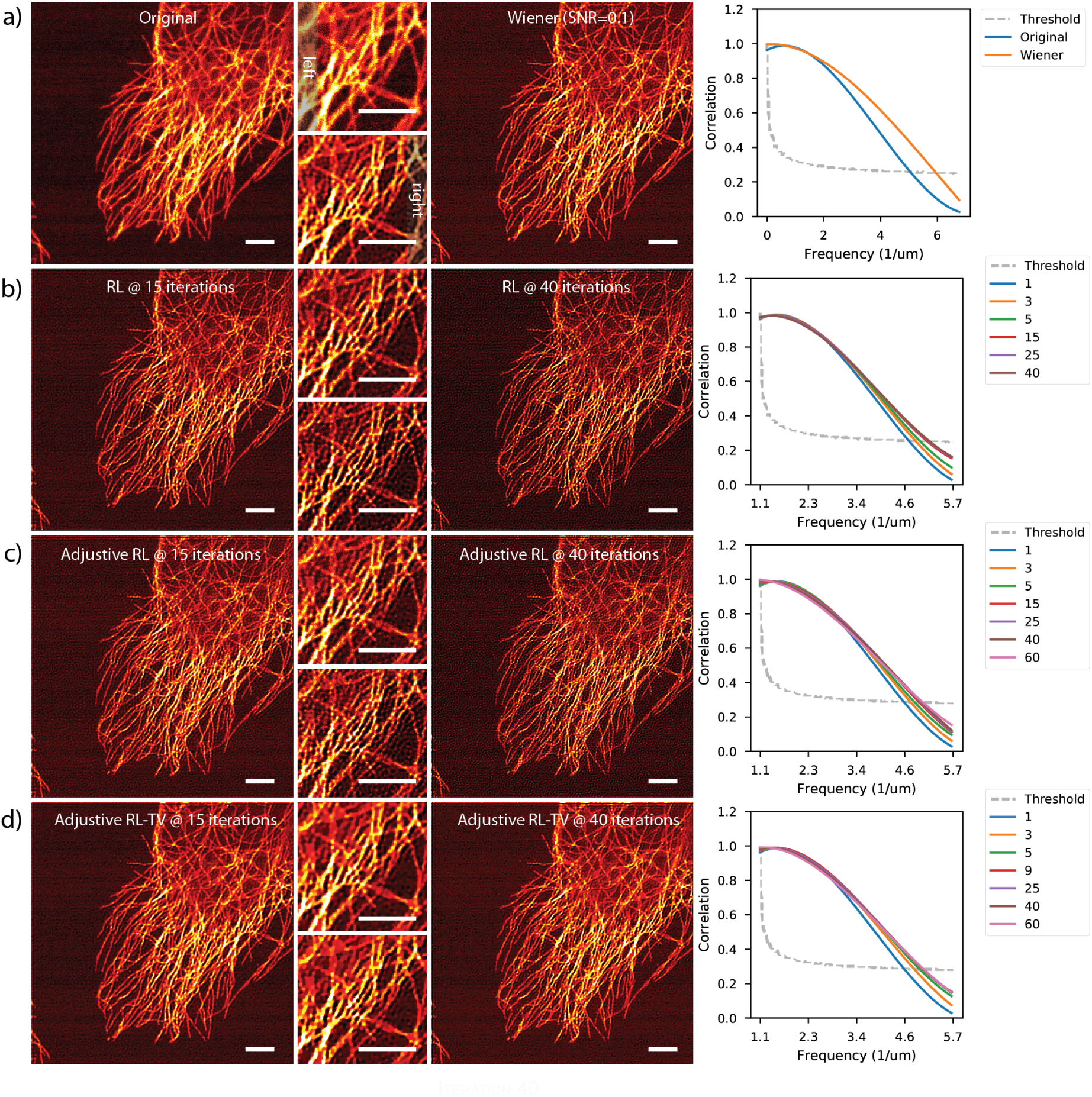
Blind Linear and Iterative Deconvolution with PSF estimated from FRC measurements. In a) results of Wiener linear deconvolution are shown. FRC measurement shows clear imporvement of effective resolution. In b) results of blind RL deconvolution, with the same image and PSF are shown. FRC measures show that the resolution does not significantly improve after 15 iterations. In c) results for Adjustive RL deconvolution are shown. In d) Adjustive RL is coupled with TV regularization. Scale bars 3*µm*.

In order to get a better understanding how our blind RL algorithms and the FRC progress metric work, we then processed several other images. In (Figure S. 6) blind RL deconvolution with TV regularizaiton is shown, with a confocal image of a vimentin stained cell (pixel size 29nm). The deconvolution converges after 39 iterations (Figure S. 6b-c)), but already after 15 iterations the result is nearly the same. The FRC measures (Figure S. 6b,d)) also show the apparent super-resolution effect (*d_min_ ≈* 100*nm*) that the iterative deconvolution is known to produce in ideal conditions [20]. In (Figure S. 7) Adjustive RL (PSF update) deconvolution results are shown for a widefield fluorescence image of a vimentin stained cell. The deconvolution effectively removes the out-of-focus haze and details are crisp after 15 iterations. The *resolution* = *f*(*iteration*) curve clearly shows (Figure S. 7b)) that the algorithm reaches maximum effective resolution after 47 iterations. In (Figure S. 9)) we compare RL deconvolution of the microtubulin stained HeLa cell image with two different PSFs (FWHM_*THEORY*_ = 170*nm*, FWHM_*FRC*_ = 200*nm*). While both of the results are rather good, the smaller PSF size leads into increase of background noise, especially at higher iteration counts, as shown in the zoomed in section in (Figure S. 9 c)). As revealed by FRC measures (Figure S. 9 b,d)), with the smaller PSF, the algorithm also converges more slowly, when compared to the blind deconvolution (Figure S. 9 a-b)). We also did two-image FRC measurements on the same images; the measurements were made by first deconvolving the two images separately with the FWHM_*THEORY*_ = 170*nm* PSF and then calculating a two-image FRC for each RL estimate pair. As shown in (Figure S. 9 e)) after 5 iterations a distortion appears in the high frequency tail of the FRC curves, which causes for the FRC values to jump to unrealistic heights.

In order to assess the robustness of the one-image FRC based progress metrics, we devised a worst case simulation of sorts, by running RL deconvolution with a small PSF (FWHM=230nm) on an image of a fluorescence layer. Such an image is more or less a worst kind of an input for a RL algorithm, as there is no useful signal to fit and thus the algorithm starts to immediately fit the noise – in fact the RL algorithm becomes rather unstable and deconvolution stops just after one iteration (Figure S. 11). With such an input the noise bump is clearly visible in one-image FRC as well, but does not affect the resolution measures or functionality of the derivatives, because of the higher FRC threshold. The appearance of the noise bump is not an effect of the bandwidth as shown in (Figure S. 6), but rather a combination of a strong constant background and a small PSF (compared to the effective resolution). Based on these observations, we also introduce the zero crossing of the second derivative of the *resolution* = *f*(*iteration*) curve as an alternative stopping point. Our reasoning is that first derivative should normally converge towards zero as the deconvolution progresses – an inflection point indicates an abnormality to this behavior. It is also possible to control the noise amplification with a smoothness constraint [13, 21, 17], or then one can try to include the background noise as an additional term term in the deconvolution algorithm [22].

To conclude, FRC is a rather valuable tool in evaluating deconvolution progress and results. Compared to two other deconvolution progress metrics that are shown for the cell image in (Figure S. 10b-c)) and fluorescence layer (Figure S. 11e-f)) it is able to provide much more information of the quality of the deconvolution results. It is possible to further increase the noise sensitivity of the one-image FRC, by turning the diagonal averaging off, as shown for the microtubulin stained HeLa cell image in (Figure S. 10a,d)); deconvolution stopping condition is reached after five iterations, just as the noise bump becomes visible in the two-image FRC.

### Working with 3D images

In fluorescence microscopy the image resolution is highly anisotropic: due to limited numerical aperture of the single objective lens microscope systems, the axial resolution (direction of the optical axis) is typically at least factor of three inferior to the lateral resolution. For this reason FSC, the simple 3D expansion of FRC, is of only limited value in fluorescence microscopy applications. In order to address this issue, we developed a FSC based method in which each Fourier Shell is divided into wedges. A single Sectioned FSC (SFSC) indexing shape consists of two such wedges that are each others mirror images (Figure S. 12 b)) – this is to take advantage of the symmetries in Fourier space. In order to calculate resolution values for the whole sphere, the dual-wedge structure is rotated in increments of *α* (Figure S. 12 a)) around an axis located on the XY plane – for each orientation a separate cross-correlation histogram and resolution value are calculated; the *α* also matches the angular size of the wedge. At *α_i_* = 2*π* the SFSC simplifies into normal FSC. As a concept the SFSC is very similar to the recently proposed Conical FSC measure [23]. They essentially differ in the way that the Fourier Sphere is indexed to produce the directional resolution measures. The SFSC is especially tuned to observe variation of resolution, when rotating around a single axis, which makes it rather fast to calculate (few sections/volume) as well as robust (large number of voxels on every section).

We compared our new SFSC measure with FRC on a STED image stack of a microtubulin stained HeLa cell. The FRC measures were made by identifying apparently in-focus planes in the 3D images and then calculating separate resolution values for the XY and XZ orientations. We also compared our SFSC measure against Fourier Plane Correlation (FPC) that was proposed for 3D image analysis in [8]. In order to facilitate the manual FRC measurement and to limit the computational load in SFSC and FPC the resolution measurements were done on a cropped 300×300×300 pixel central section of the STED image (about 1/3 of the image size). As shown in (Figure 13) the FRC and SFSC measurements are in rather good agreement: SFSC measurement at SNR_*e*_ = 0.5 threshold (one-bit), corresponds to FRC measurement at 1/7 threshold; the one-bit threshold was used in all SFSC measurements. The FPC measurement seems to work to an extent on the XY plane, but fails in all rotated directions. It is able to reveal the resolution anisotropy, but not very accurately. Also FPC seems to be very sensitive to interpolation and other artefacts in the axial direction, which does not however seem to affect the resolution measurement.

In (Figure 4) blind Wiener filtering results with two 3D images are shown; the new SFSC measure is leveraged to produce the resolution estimates, necessary for generating the PSFs. In (Figure 4a)) results obtained with a STED microscope super-resolution image are shown. The images were acquired with relatively low STED depletion intensity to ensure good contrast, and to reduce photobleaching. As it is typical to a STED microscope image, the resolution anisotropy between lateral (XY) and axial (Z) directions is considerable, as shown in the SFSC resolution plot in (Figure 4a)). The polar plot illustrates the resolution point in *µm* calculated by rotating the SFSC section at 15^◦^ increments around the Y axis. Because STED is a bandwidth un-limited technique, the power spectrum in STED images often contains very high frequencies that unfortunately, are often hidden by noise. However, the simple blind Wiener filtering as shown in (Figure 4 a)), is able to recover a surprisingly large amount of fine details. The axial haze is clearly reduced, and the effective resolution is drastically improved with previously blurred filaments, clearly visible in the results.

In (Figure 4b)) the same blind Wiener filtering approach was applied to a much larger (deep) image of Pollen recorded with a confocal microscope. Only single image was available for analysis, so the diagonal splitting was used in the SFSC calculations. The deconvolution results show dramatic improvement of contrast and details – and axial haze is effectively reduced, as indicated by the depth coloring. Even crispier details can be obtained by decreasing the Wiener regularization, but details obtained at SNR=0.005 are smooth and pleasant to view.

## Discussion

In this paper several novel blind image restoration methods were introduced that leverage FRC/FSC resolution measurements in different ways. In frequency domain de-noising methods FRC was used to find a cut-off frequency point for low-pass filtering. In deconvolution tasks, both linear (Wiener) and iterative (RL), FRC/FSC measurements were used to estimate the effective point-spread-function (PSF), directly from the image data. There are several clear benefits of estimating the PSF with FRC/FSC. Firstly, no prior knowledge – even theoretical – is needed of the microscope or the sample. Secondly, the PSF generated via FRC/FSC is always tailored to every given image. Thirdly, the PSF estimation with FRC/FSC is a single-step process, although it can also be updated iteratively if necessary. This advantage specifically made it possible to perform linear blind Wiener filtering in a very straightforward way, both in 2D and 3D. With larger images it might be of interest to divide the images into several smaller blocks [24], to adapt the PSF in the Blind deconvolution to local changes of resolution (Figure S. 14).

In iterative deconvolution (RL), FRC measures were also leveraged to observe the progress and quality of the deconvolution process. No other such metric to our knowledge exists in the literature. The *τ* and *η_k_*, as well as many other measures that can be found in the literature, mainly quantify the convergence of the deconvolution algorithm, but can not really quantitatively analyse the quality of the deconvolution results, in absence of a ground truth image. It was shown that FRC can be used to identify the deconvolution iteration at which the effective resolution is maximal, after which the algorithm starts to mainly fit noise.

**Figure 4:**
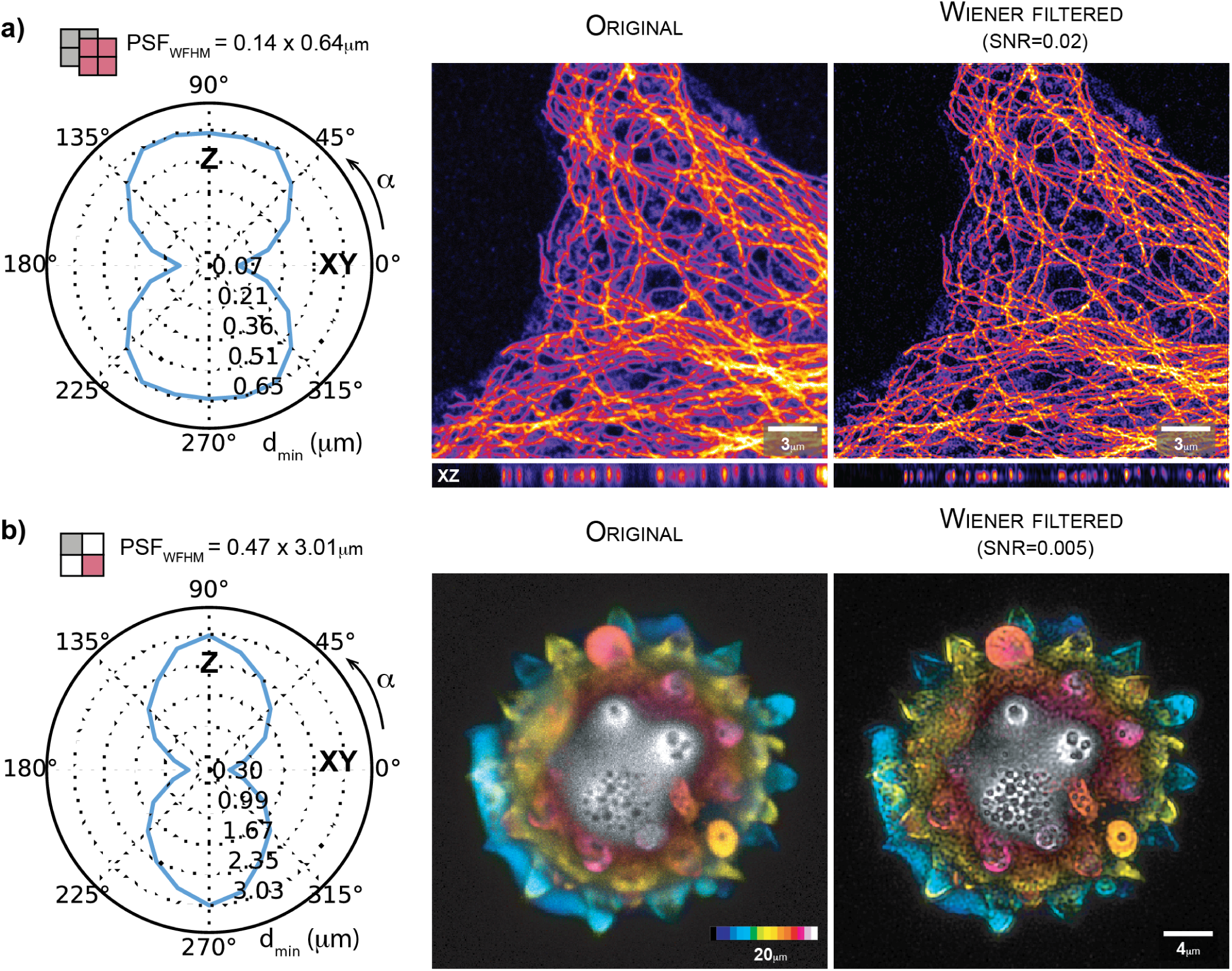
Blind Wiener Deconvolution in 3D with PSF estimated from SFSC measurements. In a), results of Wiener deconvolution of a super-resolution 3D image recorded with STED microscopy are shown. The results show dramatically improved effective resolution and reduction of axial haze. The SFSC estimate was calculated with two images. In b) Blind Wiener Deconvolution result of a confocal image of Pollen are shown. The depth coded colormap reveals dramatic reduction of axial haze, and in general contrast and effective resolution are clearly improved. The polar plots illustrate the resolution point in *µm* calculated by rotating the SFSC section at 15^*◦*^ increments around the Y axis.

In addition to new image processing methods, two major limitations in current state-of-the-art of FRC/FSC were addressed, especially with practical fluorescence microscopy applications in mind. It was shown how FRC/FSC can be calculated on single images, and how FSC can be used even in cases the optical resolution is anisotropic. In our examples, the anisotropy is evaluated in the axial direction only, but the measurement can of course be adapted for any orientation, or in the extreme case, expanded to to the full Conical FSC implementation [23] – however, with high computational cost, although Conical FSC, as well as SFSC, could to an extent be parallelised to improve speed. These methods are general, and should be of use also outside the image restoration applications.

In addition to deconvolution and image denoising that was the focus of this paper, there are also several other image processing/analysis tasks that FRC/SFSC could be applied to. In (Figure S. 16) we entertain the idea of combining FRC/FSC with other image quality assessment parameters, to produce quantitative measures of image quality for e.g. high-content screening applications; a similar method was recently proposed for assessing the quality of localization super-resolution microscopy image reconstructions [25]. FRC is also very sensitive to the focal position: as shown in (Figure S. 15) FRC actually behaves rather similarly as several autofocus metrics, which might be an interesting future application – and of course, a good way to identify an in-focus plane at post-processing stage. Our Open Source MIPLIB software library that was used to perform all the demonstrated image analysis and processing tasks, may help in such future applications.

## Methods

### One and two-image resolution measurements with 2D images

One- and two-image FRC measurements were both done according to

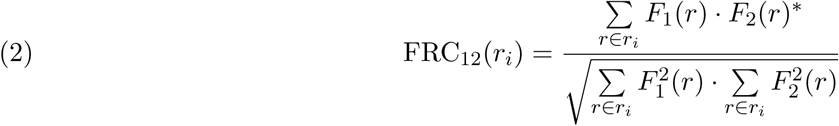

, where *F*_1_ and *F*_2_ are the Fourier transforms of the two images and *r_i_* the *i*th frequency bin. Prior to the FRC calculations, in two-image case, the two images were registered by a phase-correlation based method [26]; in single-image case the splitting was performed to produce the four sub-images. A Hamming window was applied to each image to suppress edge effects and other spurious correlations. In all our single-image FRC measurements the diagonal splitting pattern (Figure 1a)) was used, if not otherwise specified. In two-image FRC the 1/7 resolution threshold [8, 7] was used to determine the numerical resolution value. The one-image threshold SNR_*e*_ = 0.25 is based on information theory. The SNR_*e*_ = 0.25 threshold curves introduced in [10] reads

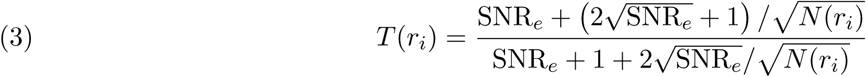

, where *T* (*r_i_*) is the threshold value, *N* (*r_i_*) the number of pixels/voxels at Fourier ring/shell *i*, and SNR_*e*_ the expected SNR value at the cut-off point. The SNR_*e*_ threshold was selected experimentally, to very closely match the numerical values obtained with two-image FRC measures at 1/7 threshold; it is nearly equivalent to the *half-bit* curve proposed in [10] (a bit higher).

### Resolution measurements with 3D images

Measurements on 3D images were done with our Sectioned Fourier Shell Correlation measure (SFSC)

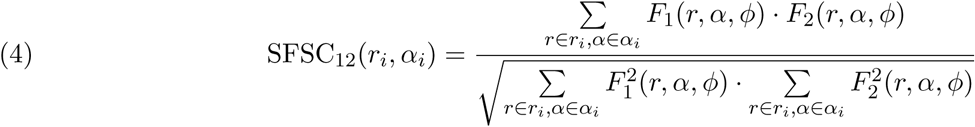

, where *F*_1_(*r, α, φ*) and *F*_2_(*r, α, φ*) denote the voxels in two Fourier transformed images that are located (I) at a given distance *r_i_* from the origin and (II) within an orientation sector, defined by *α* and *φ*. We compared the SFSC measures against FPC [8] as well as FRC measures.

The SFSC/FPC/FRC analyses with the 3D images were performed with the same logic as the FRC measures on the 2D images. Before data analysis each 3D image was resampled to isotropic spacing using linear interpolation. Single image splitting was achieved with the diagonal splitting (Figure 1a)); only one diagonal was used to limit the computational effort in SFSC/FPC. In the axial direction two consecutive axial (z) layers were simply added together, to maintain the image proportions, and to not introduce additional offsets. Threshold curve based on SNR_*e*_ = 0.5 (*one-bit*) was used with SFSC, whereas with both FPC and FRC 1/7 threhold produced more reasonable numerical values.

Because of the anisotropic sampling that is typical to 3D fluorescence microscopy images, the frequency axes in SFSC and FRC curves need to be corrected to compensate for it. In this paper this was achieved by multiplying the image pixel/voxel size by factor *k*(*θ*):

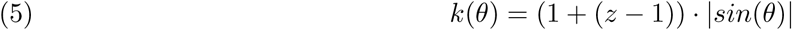

, where *z* is the sampling anisotropy factor (e.g. 2 for an image sampled with half the sampling rate in depth (Z) with respect to the lateral (XY) direction) and *θ* denotes the rotation angle with respect to the XY plane, which in case of SFSC is a multiple of *α*. If no such correction is made, all the numerical values calculated with FRC/SFSC at orientations *θ* ≠ 0 + *nπ* will have unrealistically high numerical resolution values. Also FPC overestimates the resolution values on every rotated plane, but the same simple correction does not seem to work.

### Frequency Domain Low-Pass Filtering

The frequency domain filtering was performed by first estimating the effective image resolution with FRC/FSC, and then using it as a cut-off frequency for a low-pass Fourier domain filter. Three different types of Fourier space filters were used in this work: (I) an ideal low-pass filter (ILPF), (II) a Butterworth low-pass filter and (III) a Gaussian low-pass filter.

An ILPF can be defined as:

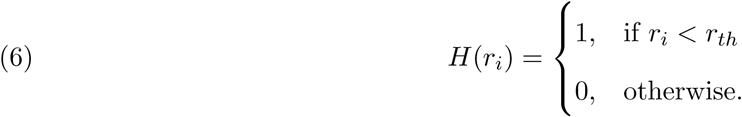

, where *r_i_* is a polar distance from the center of the Frequency domain filter (zero frequency) and *r_th_* is the distance at the cut-off frequency, obtained with FRC/FSC. Frequencies after the cut-off are simply clipped to zero.

A Butterworth low-pass filter (BLPF) on the other hand is defines as:

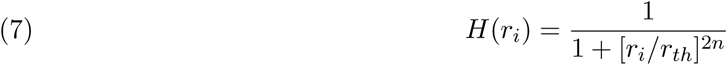

, where *n* denotes the degree of the filter, which controls how sharply the transition from pass-band (allowed frequencies) to stop-band (filtered frequencies) is made. When compared to ILPF, BLPF has nearly equally flat (unity) response in the pass-band, but transitions to stop-band more smoothly, thus avoiding a strong discontinuity at the cut-off; at the *r_th_ H*(*r_i_*) = 0.5.

A Gaussian low-pass filter (GLPF) is defined as:

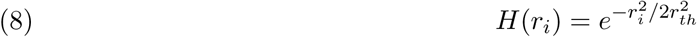

Similarly to BLPF, the GLPF has a smooth transition from pass-band to stop-band. The Gaussian function however, is not flat in the pass-band and transitions more slowly from pass-band to stop-band, which means that it may in some cases blur the filtered images and perform less than optimally, when filtering out noise. At the *r_th_ H*(*r_i_*) = 0.607.

### Image Deconvolution

Image formation in a microscope can be described as a convolution of every object sample point with the PSF:

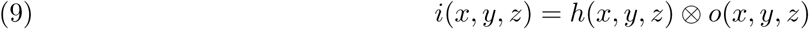

, where *i*(*x, y, z*), *h*(*x, y, z*) and *o*(*x, y, z*) are the measured image, the PSF and the original sample object, respectively. By image deconvolution [27] one attempts to revert the blurring effect of the microscope, and thus increase image contrast and effective resolution, by using the PSF as a prior information. Deconvolution can be performed in a single step, e.g. by Wiener or Tikhonov filtering [28] – or then iteratively, e.g. by Richardson-Lucy (RL) [29, 30]. In this paper we use the Wiener and RL algorithms.

The Wiener deconvolution algorithm is based on inverse filtering, and takes advantage of the fact that a convolution operation in the spatial domain, becomes a multiplication in the frequency domain – and thus a relatively simple single-step deconvolution can be realised as follows:

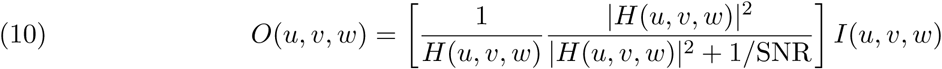

, where *O*(*u, v, w*), *I*(*u, v, w*), *H*(*u, v, w*) are the Fourier space representations of the estimate for the original object, the observed image and the PSF. *|H*(*u, v, w*)|^2^ is the power spectrum of the PSF. The regularization term 1/SNR can also be written as *|N* (*u, v, w*)|^2^*/|O*(*u, v, w*)|^2^, where *|N* (*u, v, w*)|^2^ is the power spectrum of the noise and *|O*(*u, v, w*)|^2^ is the power spectrum of the original object; neither of the two terms are known, which means that usually the value is decided on case-by-case basis, based on the subjective quality of the deconvolution results. The regularization factor is weighted by the power-spectrum of the PSF; it will have a higher effect at high frequencies, as the power-spectrum approaches zero value.

The iterative Richardson-Lucy (RL) algorithm can be described by

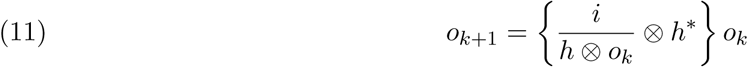

, where *o_k_* and *o_k_*_+1_ are the current and next object estimates, i is the original image, *h* is the PSF and *h^∗^* its mirrored version. In the equation, the pixel indexes have been omitted to allow a simple presentation of the algorithm.

With both Wiener filtering and RL, PSF estimate was generated on the basis of an FRC measurement on the original image data: the FRC resolution value was simply used as an FWHM value for a Gaussian PSF. With 3D images separate values were used for lateral and axial directions. In the *Adaptive RL* algorithm we updated the PSF during RL iteration, in which case a FRC measure was taken after each iteration step and a new PSF was generated according to the new FWHM width.

In addition to updating the PSF estimates (when so desired) the FRC measures of the intermediate estimates were used to observe the progress of the RL deconvolution: the deconvolution was considered fully converged, when effective resolution reached its maximum value. We also defined an alternative threshold for stopping the iteration, based on the rate of change of the effective resolution (1*µm/it*) in our example. We compared our metric against two previously published ones *τ*_1_ and *η_k_* [31, 32, 33].

With *tau*_1_ the relative difference between two subsequent deconvolution estimates is measured:

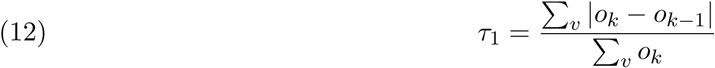

, where *o_k_* is the current estimate and *o_k−_*_1_ the previous one. *η_k_* is a measure of convergence,

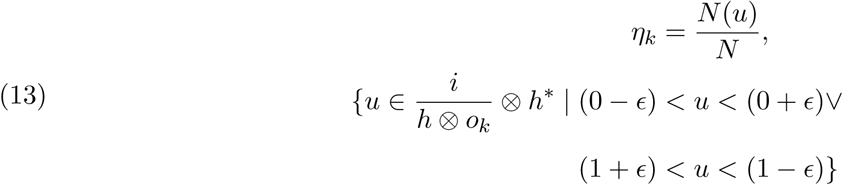

, where *N* is the total number of pixels, *N* (*u*) is the number of pixels that are not currently converging and ϵ is the convergence epsilon, 0.05 in results shown this paper.

Total Variation regularization [17] with *λ_T V_* = *⋅*10^*−*4^ was used as a smoothness constraint in (Figure 3 d)) as well as (Figure S. 6), elsewhere unregularized RL algorithm was used.

## Code Availability

The FRC/FSC measurement functions as well as all the image processing, analysis and data visualization tools used in this paper are available as Open-Source Python software library, called *Microscope Image Processing Library* (MIPLIB)(Note S. 3).

## Test Images

The test images consist of various types of confocal as well as STED microscope images that were acquired with a variety of commercial and custom-built microscopes. None of the samples were specifically prepared for this paper, but a short description for each is given in (Note S. 2).

## Data availability

The data that supports the findings of this study are available from the corresponding author upon request.

## Abbreviations

FRC: Fourier-ring-correlation
FSC: Fourier-shell-correlation
STED: stimulated emission depletion
SFSC: Sectioned FSC
FWHM: full-width-at-half-maximum
RL: Richardson-Lucy
WF: Wiener filter
SNR: signal-to-noise ratio

## Acknowledgments

The authors thank Dr. Paolo Bianchini and Prof. Colin J. R. Sheppard (Istituto Italiano di Tecnologia) for useful discussion, and Elena Tcarenkova (University of Turku) for help with obtaining the Abberior STED images.

## Author contributions

S.K., G.T., M.C., and G.V. conceived the idea. S.K. and G.V. planned the studies and the experiments. A.D. and G.V. supervised the project. S.K. wrote the software and perform the majority of the experiments. T.D. performed the three-dimensional confocal experiments. S.K. and G.V. analyses the data. S.K., G.T., M.C., T.D., A.D. and G.V participated in extensive discussions during the course of the research. S.K. and G.V. wrote the manuscript. G.T., M.C., T.D., and A.D. participated in revising the manuscript.

## Supporting Information

**Figure S. 1.**
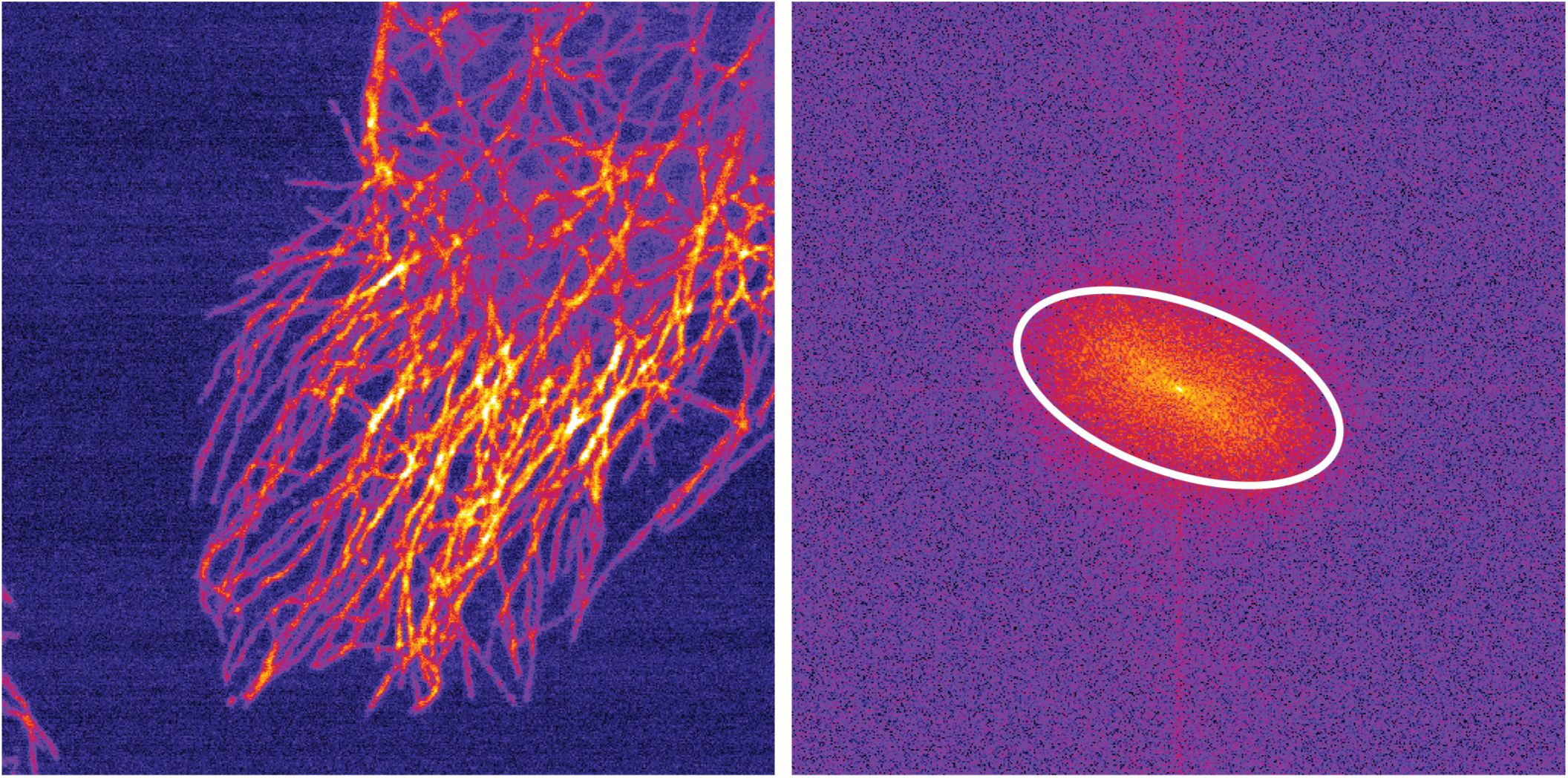
Regarding FRC and special symmetries. As was shown in (Figure S. 5) the diagonal single image splitting is mostly invariant of the splitting direction, except for in cases such as the microtubulin stained HeLa cell image here, the features in an image are mainly oriented to a certain direction, which produces a very anisotropic power spectrum.

**Figure S. 2.**
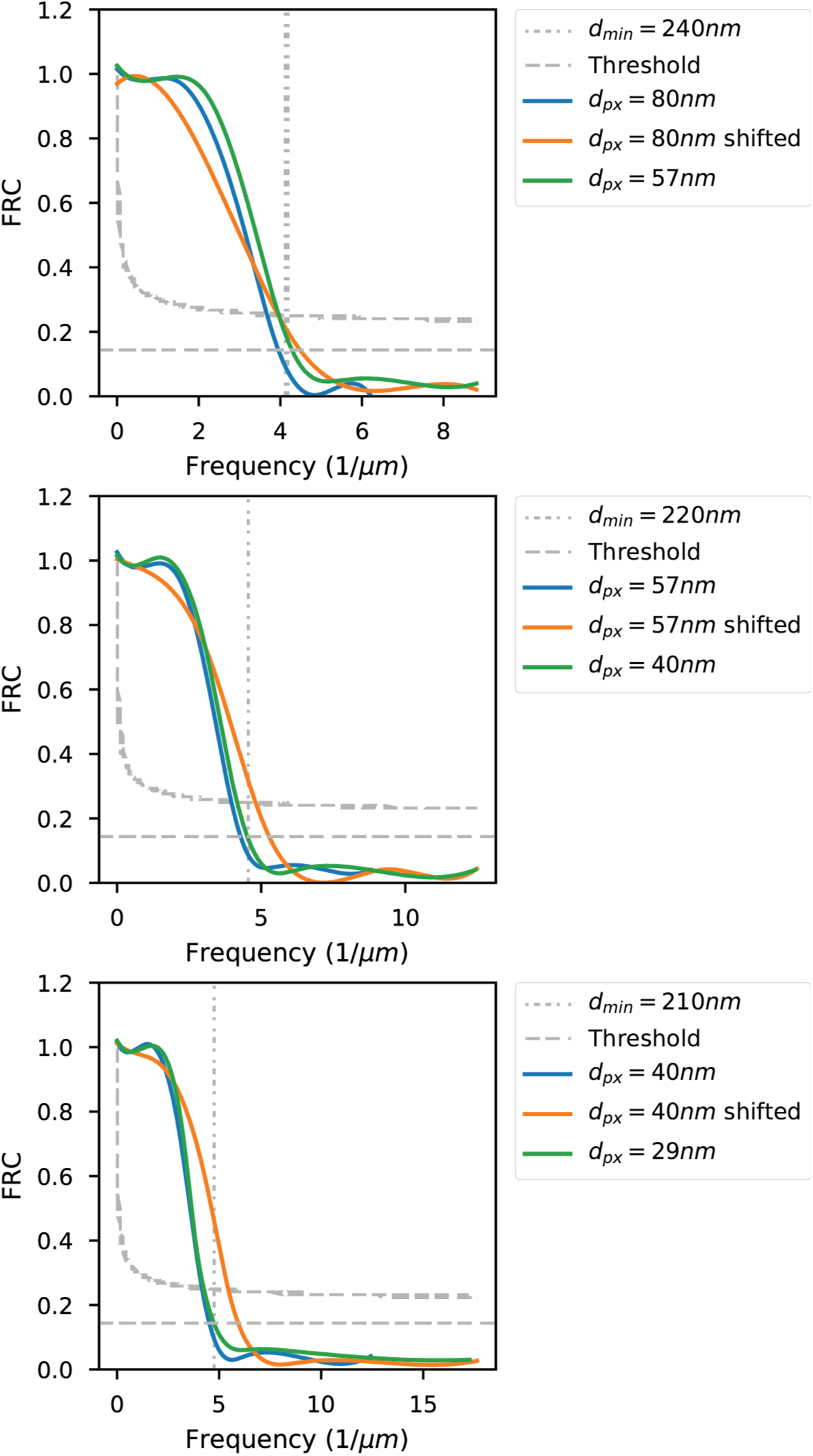
Simulating the effect of 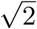 shift in two image FRC. The frequency axis offset effect is simulated in a regular dual-image FRC measurement, with the confocal dataset of vimentin stained cells, with different sampling densities. The shifted result was obtained by first applying the shift to one of the two images, after which FRC was calculated, and then the frequency axis was rescaled by the 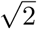 factor. According tot the theory (Note S. 1), FRC after the shifting should be somehow similar to that of an image with 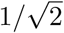 smaller pixel size. Surprisingly, this is rather exactly the case in the first example (80nm, 57nm pixel sizes). As with one-image FRC, also in this two image simulation the shift compensate FRC curve appears to slightly overestimate the resolution at smaller pixel sizes; however, even at worst, with the (40nm/29nm) pixel size the difference is only circa 25nm. The SNR=0.25 threshold correlates well with the 1/7 threshold of the regular FRC, as in the one-image case.

**Figure S. 3.**
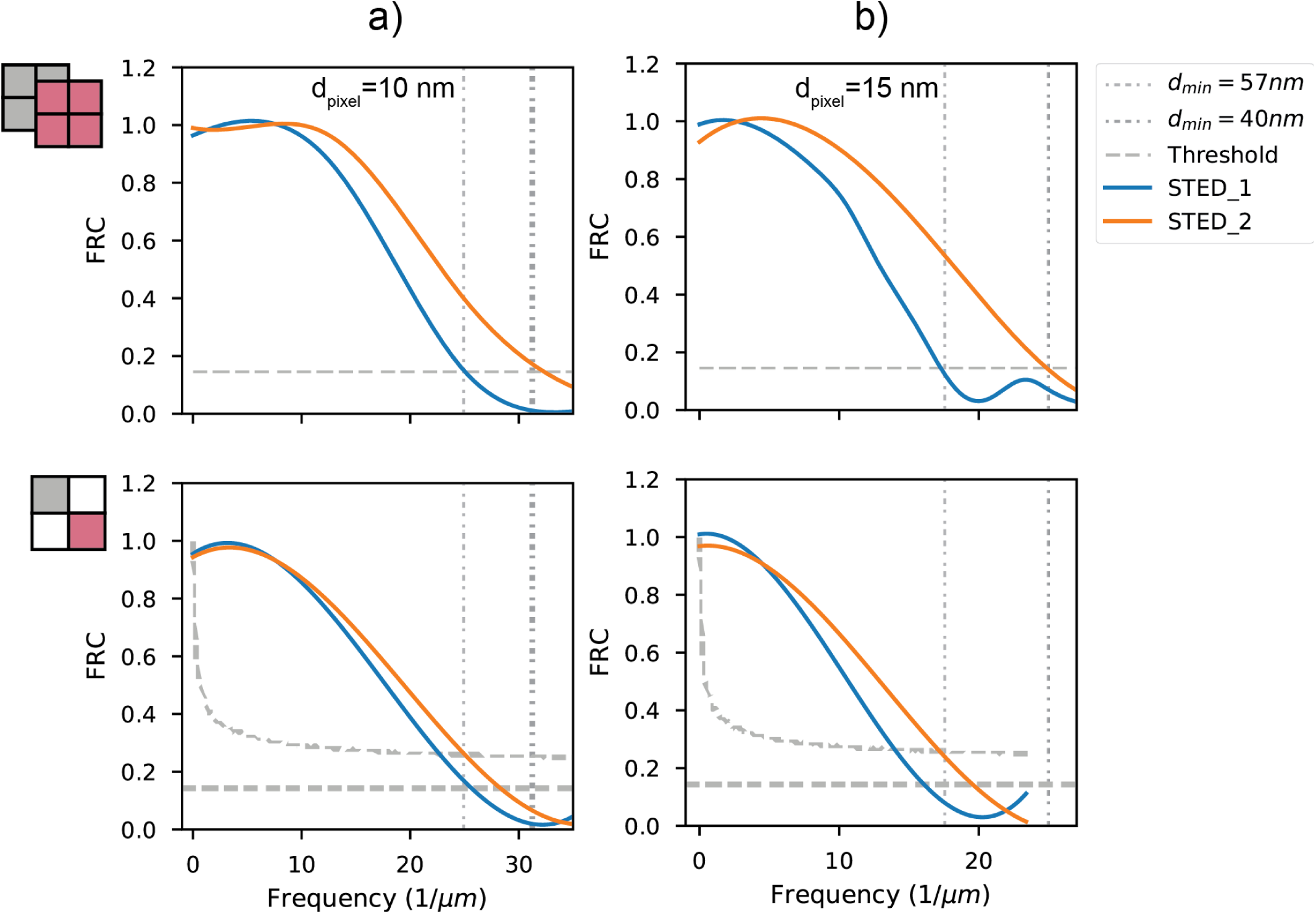
Investigating the high-frequency behavior of single image splitting with STED images of Crimson nanoparticles. One- and two-image FRC measures of two sets of STED super-resolution images (with STED 1 and without STED 2 time gating [1]) of 40nm Crimson fluorescent nanoparticles with 10nm/15nm pixel size are shown. It is evident that at the high frequencies the SNR=0.25 somewhat underestimates the resolution. At the 1/7 threshold on-e and two-image FRC produces practically identical results with the non-gated images, whereas with the time-gated images the one-image FRC is somewhat attenuated, as the resolution approaches the Nyquist limit; the difference however is only 10nm in both a)(30nm *→* 40nm), and b) (40nm *→* 50nm).

**Figure S. 4.**
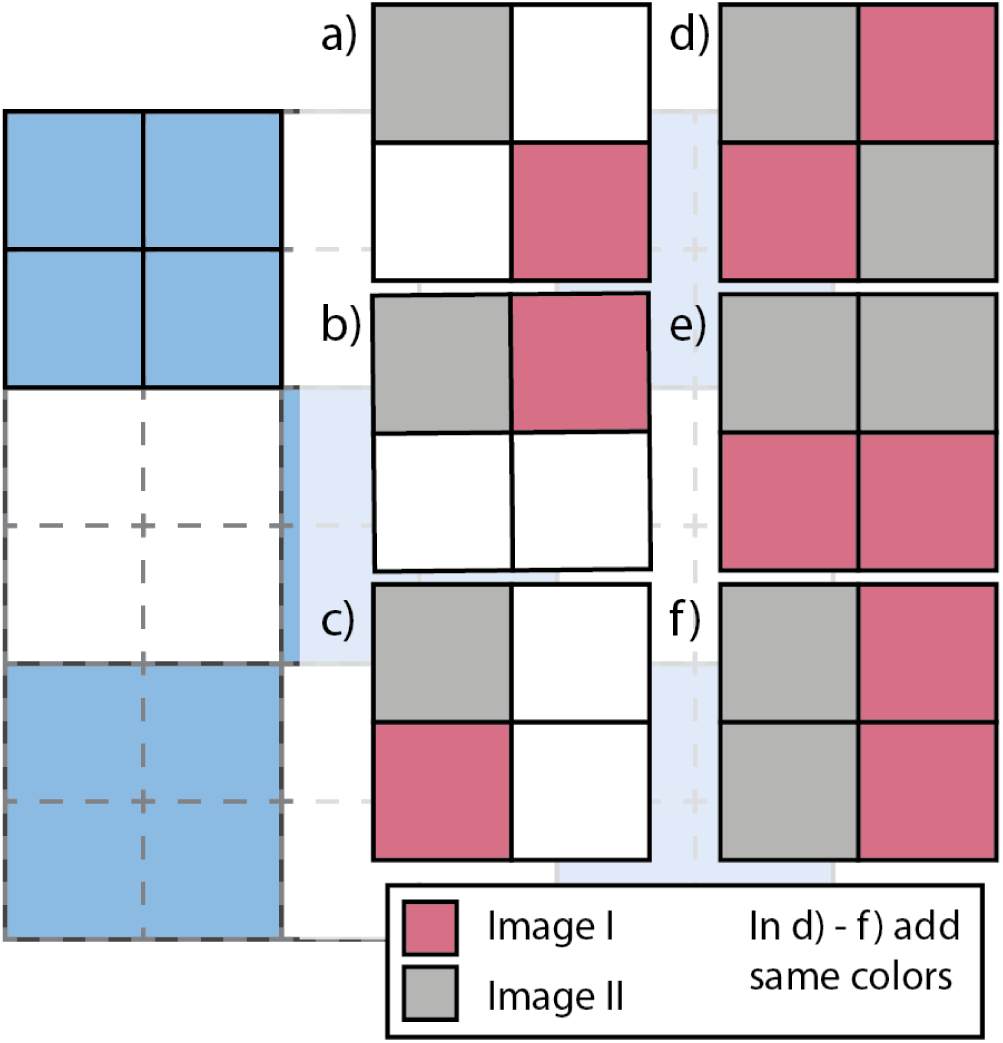
Methods to split a single image into two sub-images are illustrated. In order to perform the splitting two pixels need to be created from every four pixel group in the original image. In a)-c) the splitting is done by sub-sampling; two pixels in every four pixel group are selected to form the two sub-images. In d)-f) two pixels are added together in every four pixel group to create a new pixel value for a sub-image. The alternative a) can be realized as shown in the figure or by using the unselected pixels instead.

**Figure S. 5.**
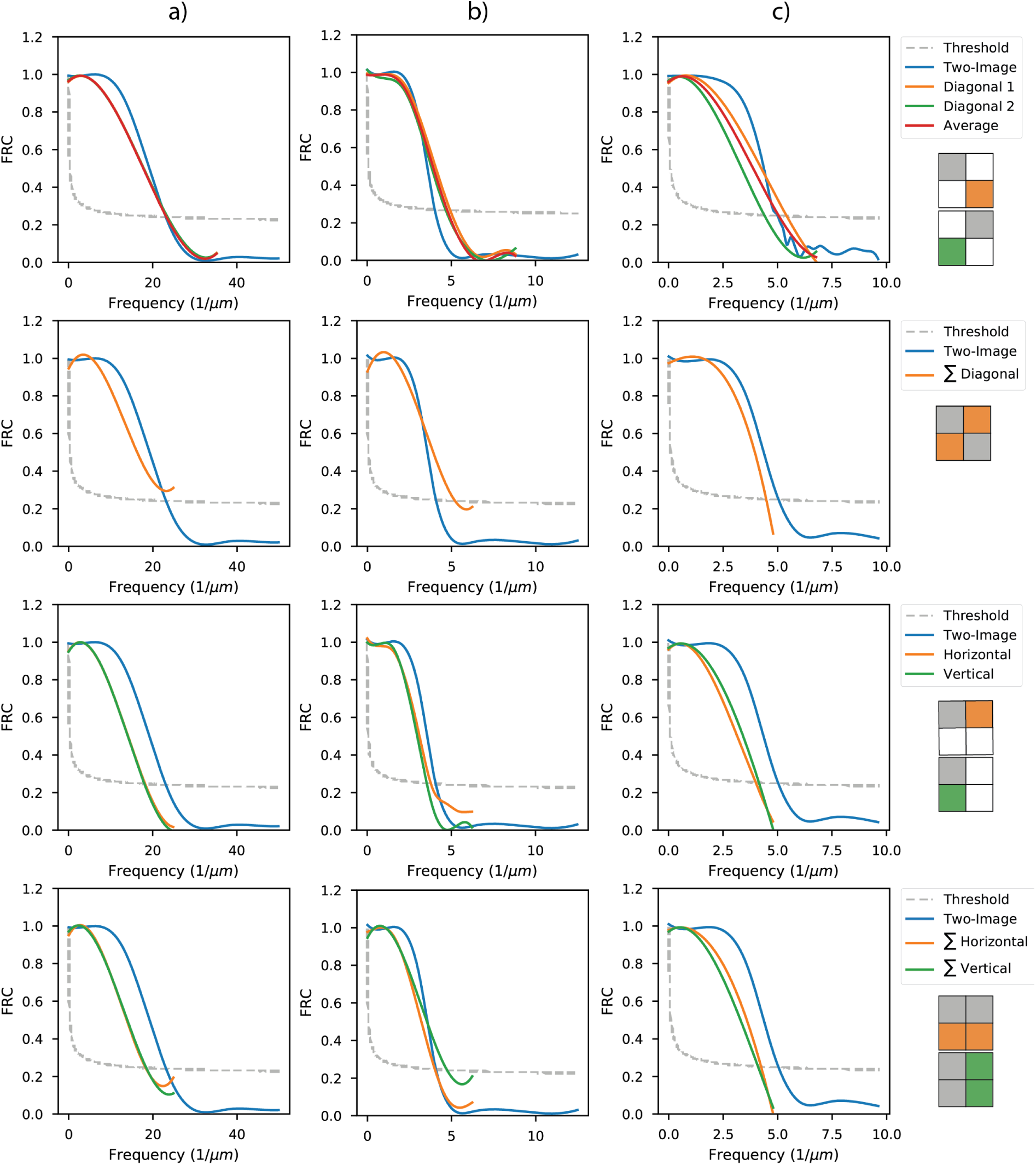
Comparing possible single image splitting schemes. In a) results obtained with a STED super-resolution image of Crimson fluorescent beads (10nm pixels size) are shown. Only the diagonal splitting produces accurate results. In b), results obtained with a fixed cell confocal image with vimentin staining are shown (40mm pixel size). Again, the diagonal splitting produces excellent results, with no variation between the two diagonals. Despite apparent over estimation, the one-image FRC value at the SNR=0.25 is practically equal to two-image FRC at 1/7 threshold. Interestingly the diagonal summing scheme (second row) produces a similar curve; the difference does not thus appear to be due to the rescaling. In c) results obtained with microtubulin stained fixed cell image (main image used in the article, 52nm pixel size) are shown. Only the diagonal splitting works properly again. The two diagonals now produce different results, because the image has a strongly asymmetric FFT (Figure S. 1).

**Figure S. 6.**
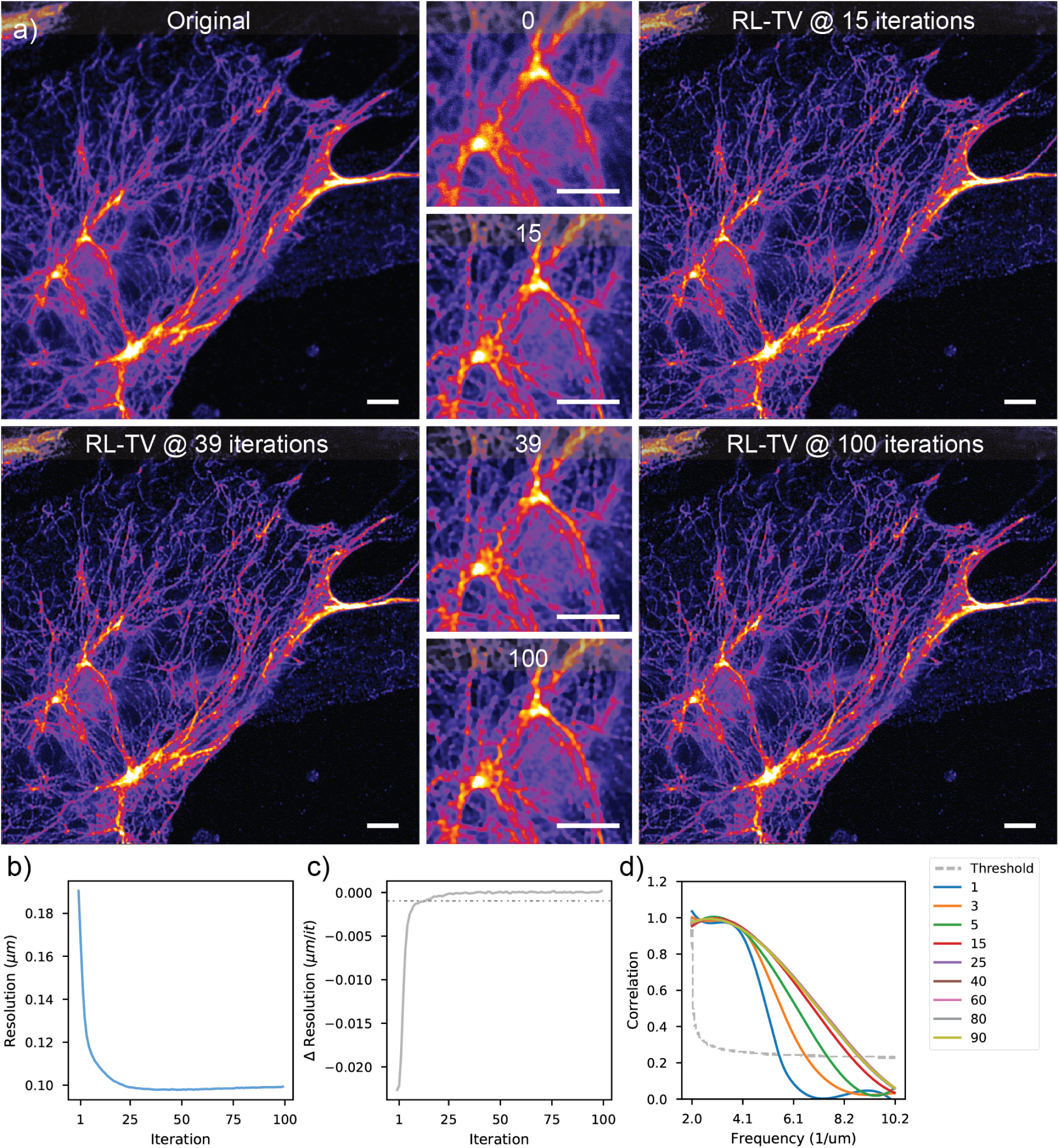
Blind RL-TV deconvolution of a confocal image of a Vimentin stained cell. The effect of TV regularization (*λ* = 5 *⋅* 10^*−*4^) is demonstrated with a confocal image of a fixed vimentin stained cell (29nm pixel size). As shown in b-c) the deconvolution converges after circa 30 iterations (max. resolution point 39 iterations) and the result remains remarkably stable henceforth. An apparent super-resolution effect is shown as the resolution peaks at approx. 100nm. The FRC measures in d) show an absence of any noise “bumps”. Scale bars 3*µm*.

**Figure S. 7.**
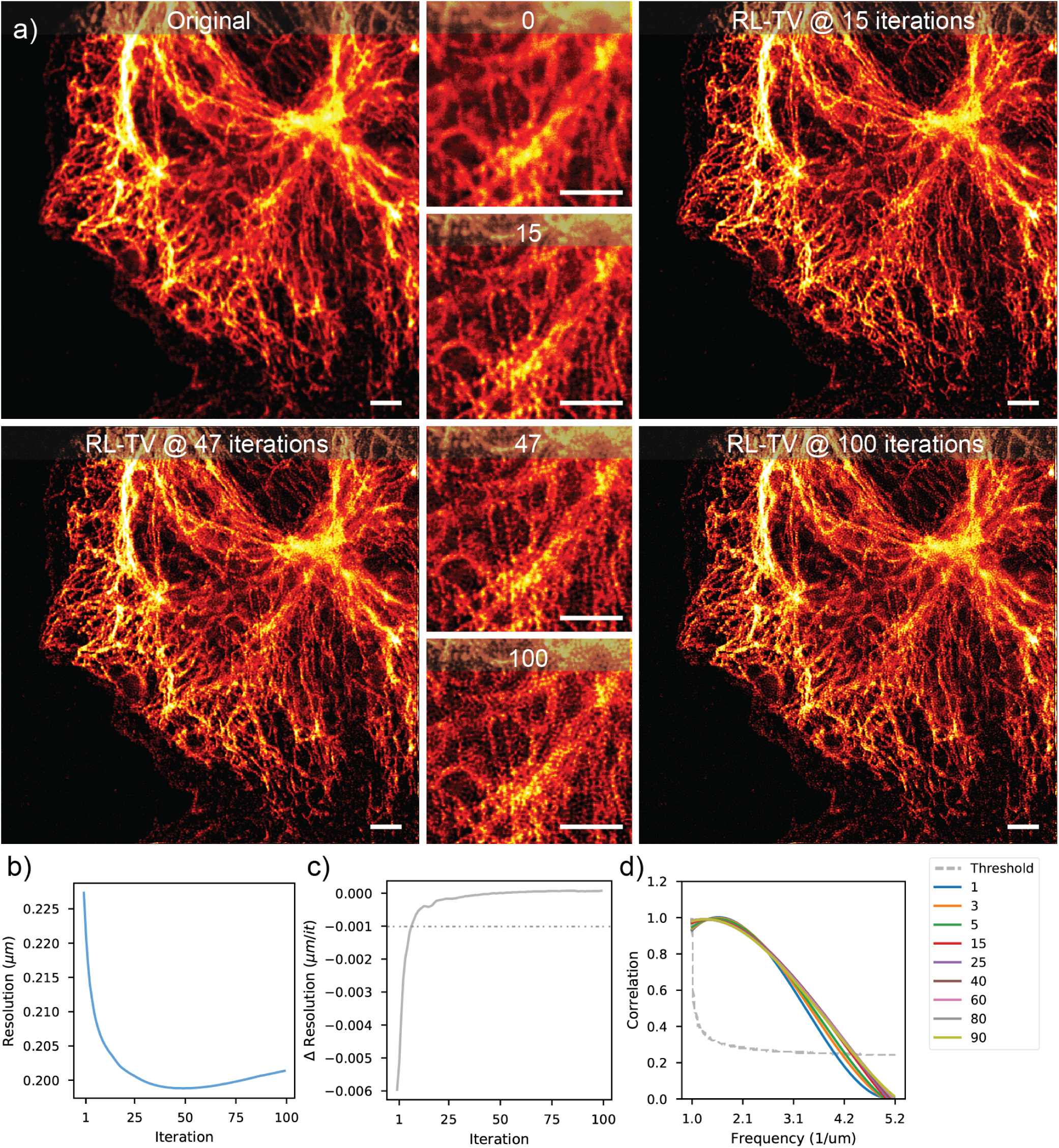
Blind Adjustive RL deconvolution of a widefield image of a Vimentin stained cell. Adjustive blind RL deconvolution (with PSF update at every step) is demonstrated with a widefield image of a vimentin stained fixed cell (pixel size 56nm). As show in b-c), the deconvolution converges at 47 iterations, after which the result starts to deteriorate due to pixelation (no regularization). Only modest resolution gain is obtained b,d), but all of the out-of-focus blur is removed. Scale bars 3*µm*

**Figure S. 8.**
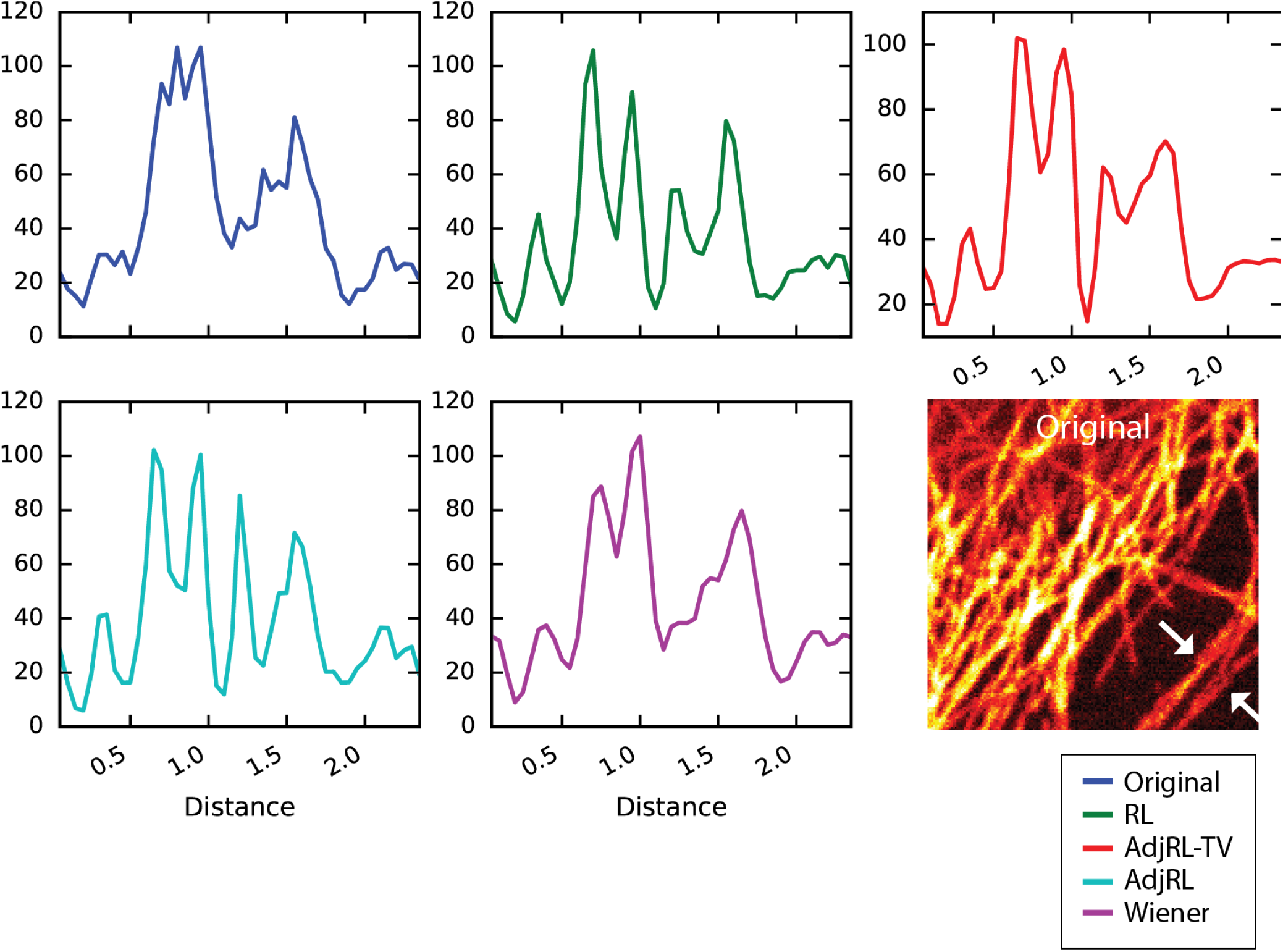
Comparison of deconvolution results based on line profile measures. The different deconvolution algorithms are compared by line profile measurements on the microtubulin stained HeLa cell sample. The results are nearly identical, as was also suggested by FRC.

**Figure S. 9.**
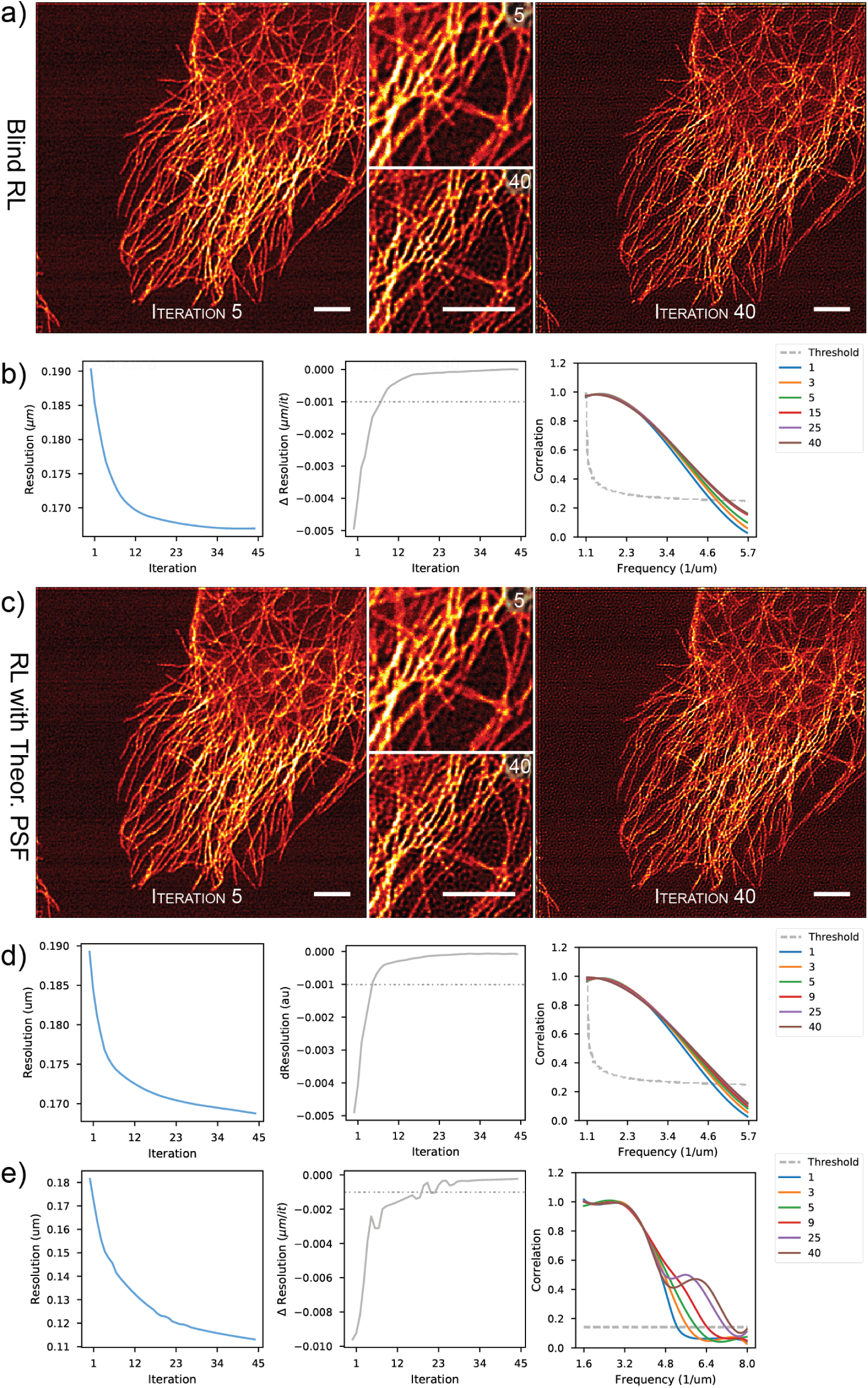
Progress of the blind RL deconvolution is observed with FRC measurements. In a) results of the blind RL deconvolution after 5 and 40 iterations are shown. As shown in b) the maximum resolution (zero crossing of ∆Resolution) is reached after 38 iterations, but only minor improvement is made after approx. 10 iterations. In c) RL deconvolution results with the same image and a smaller (FWHM = 170nm vs. 200nm) PSF are shown. Because of the smaller PSF, the noise is much more apparent after 40 iterations. One-image FRC based plots in d) show that the deconvolution converges more slowly. In e) the RL deconvoluiton with 170nm PSF is observed with two image FRC. Unrealistically high resolution values are produced, because the FRC measurement is distorted by a strong noise peak that becomes visible after five deconvolution iterations in FRC. In b,c,f) the dotted horizontal line denotes an alternative deconvolution stopping position (∆Resolution *≤* 1nm/it).

**Figure S. 10.**
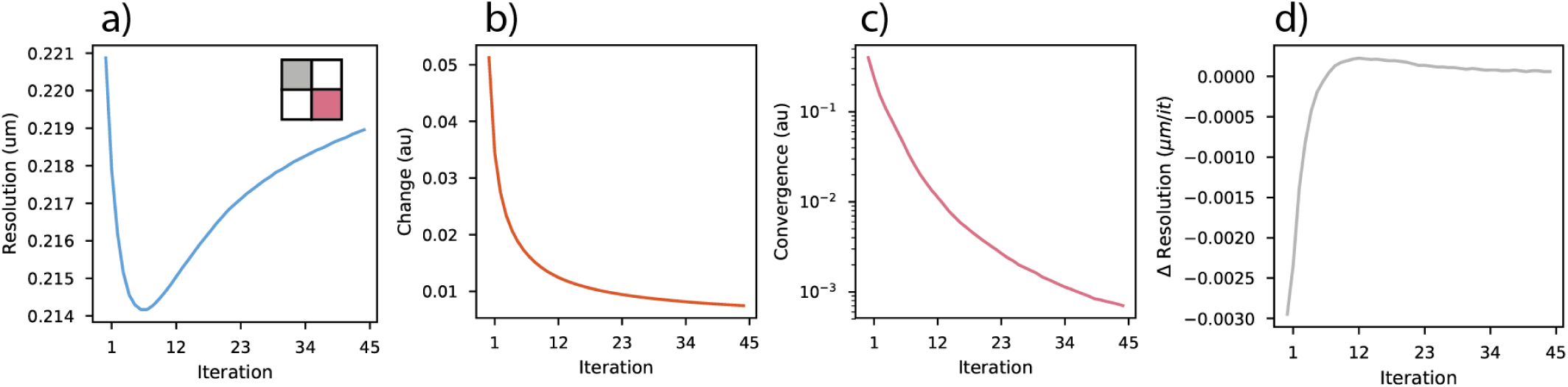
Additional measures of the RL deconvolution progress with the microtubulin stained HeLa cell. In a) on image FRC measure without the diagonal averaging is shown, for the deconvolution process with the theoretical PSF (main text, Figure 4). The measure is very sensitive to noise and the first derivative d) of a) reaches zero already after 5 iterations. The alternative progress measures *τ*_1_ b) and *η_k_* c) are rather useless.

**Figure S. 11.**
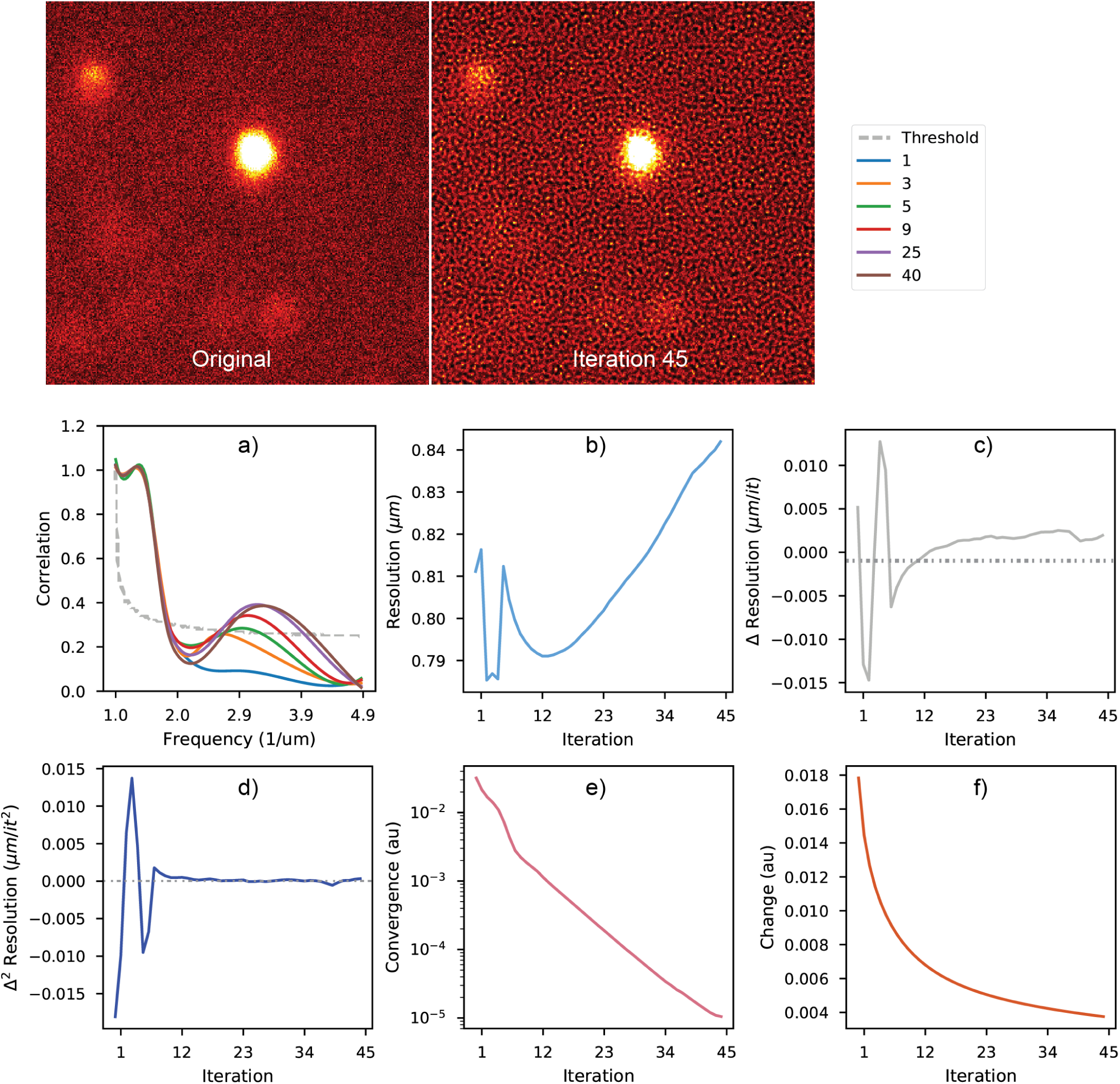
RL deconvolution of a fluorescence layer. RL deconvolution was run on a fluorescence layer (with some fluorophore aggregates visible). The one-image FRC measures a) show a noise bump starting to form, right from the first iteration. The resolution/iteration plot shows that the deconvolution is rather unstable – after some oscillation the effective resolution starts to decrease as a function of the iteration count. The first c) and second d) derivatives of b) cross zero after one iteration, indicating that the deconvolution should stop. Two other deconvolution progress parameters *η_k_* e) and *τ*_1_ f) do not work.

**Figure S. 12.**
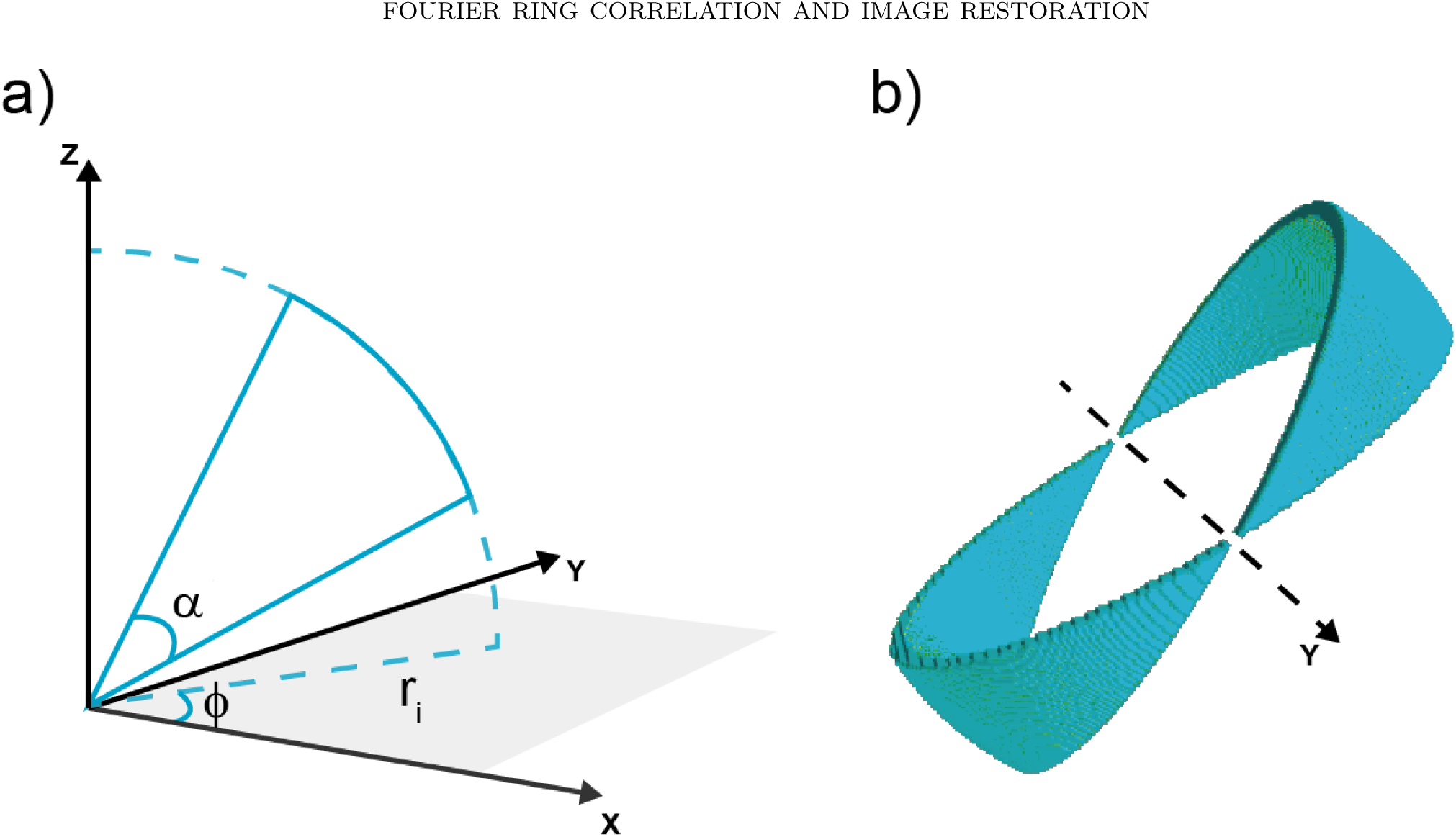
Illustrating the principle of SFSC Fourier Shell Indexing. In a) the indexing parameters in 3D coordinate system are explained. *α* (elevation), defines the thickness of the section and the orientation of the shell in relation to XY plane; the orientation is a multiple of *α* and is changed as a part of the iteration process. *φ* (azimuth) defines the rotation axis on the XY plane. *r_i_* is the radius of the *i*th shell. In b) volume rendering of a single indexing structure is shown; the rotation in this case is done around the Y axis.

**Figure S. 13.**
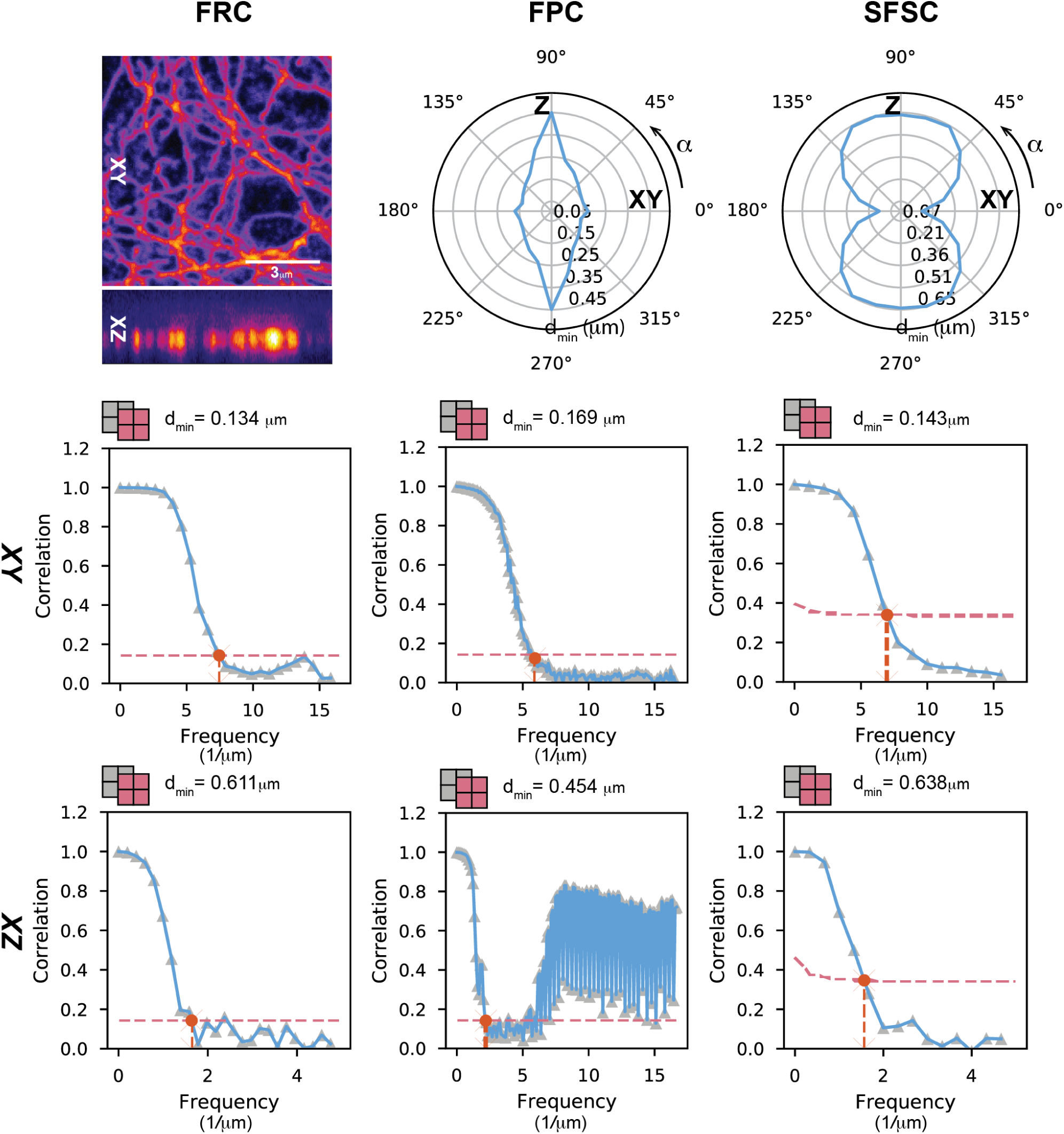
3D resolution measurements with FRC, SFSC and FPC. The resolution measures with FRC, FPC and SFSC are compared on the STED image stack, by resolution measures in XY and XZ directions. In addition, for FPC and SFSC 360 degree polar plots are shown. FRC and SFSC are in good agreement and produce reasonable resolution values. FPC seems to work relatively fine in XY but fails on any rotated plane, as is evident from the polar plot representation. FPC also seems to be very sensistive to (interpolation) artefacts.

**Figure S. 14.**
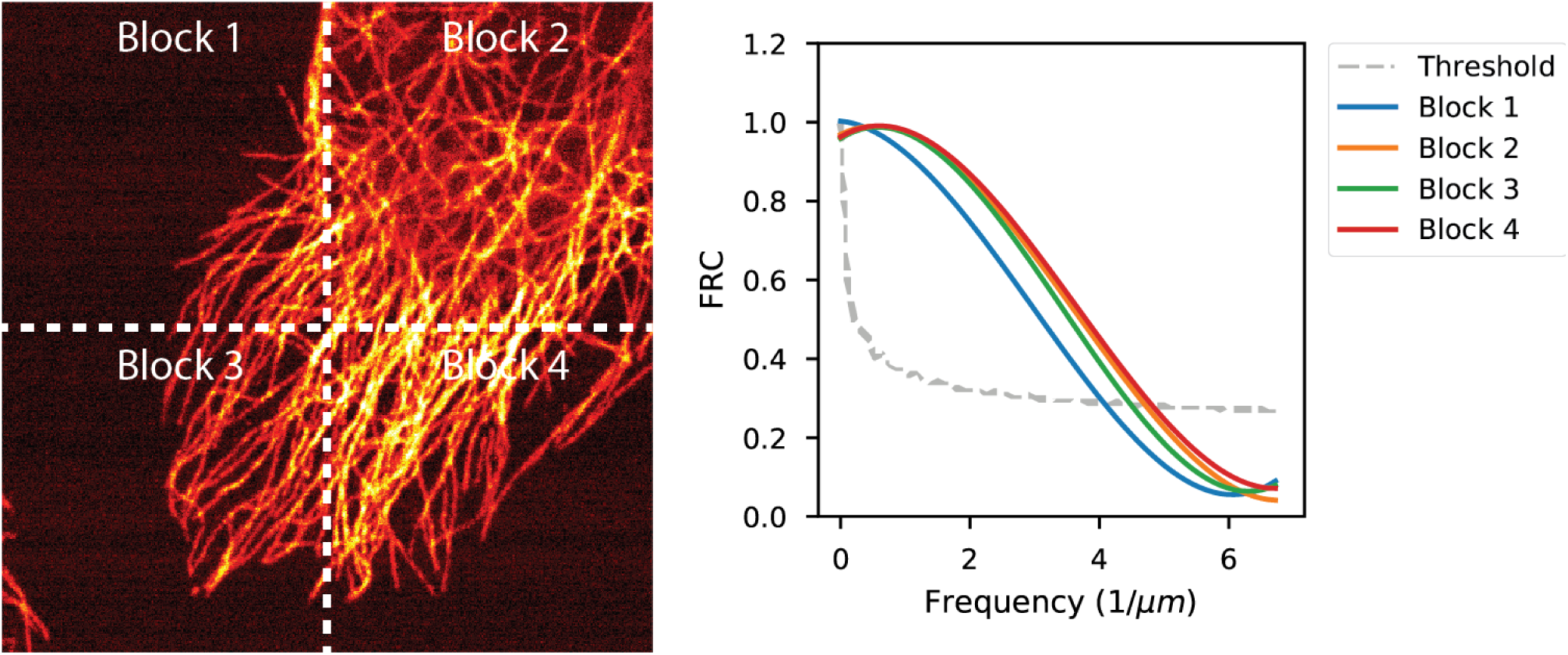
Block based FRC. It is demonstrated with the microtubulin stained HeLa cell that it is possible to easily calculate local resolution values with the one-image FRC.

**Figure S. 15.**
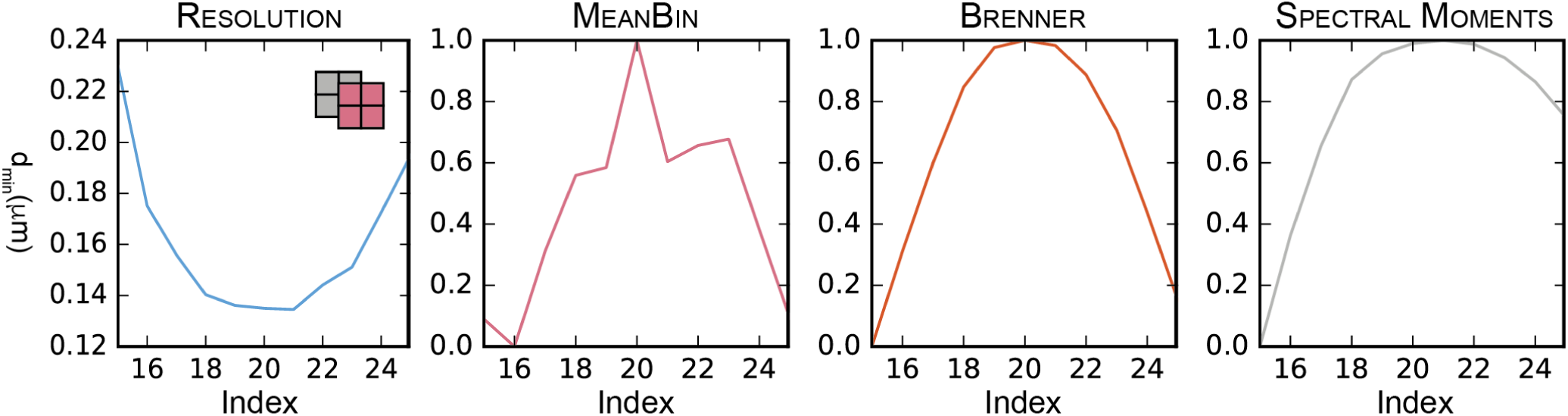
FRC dependency on focal position in a 3D stack. The resolution measured with FRC at different axial positions in the STED image stack (main article **Figure 5**), is shown and compared to several auto-focus metrics; see [2] for details.

**Figure S. 16.**
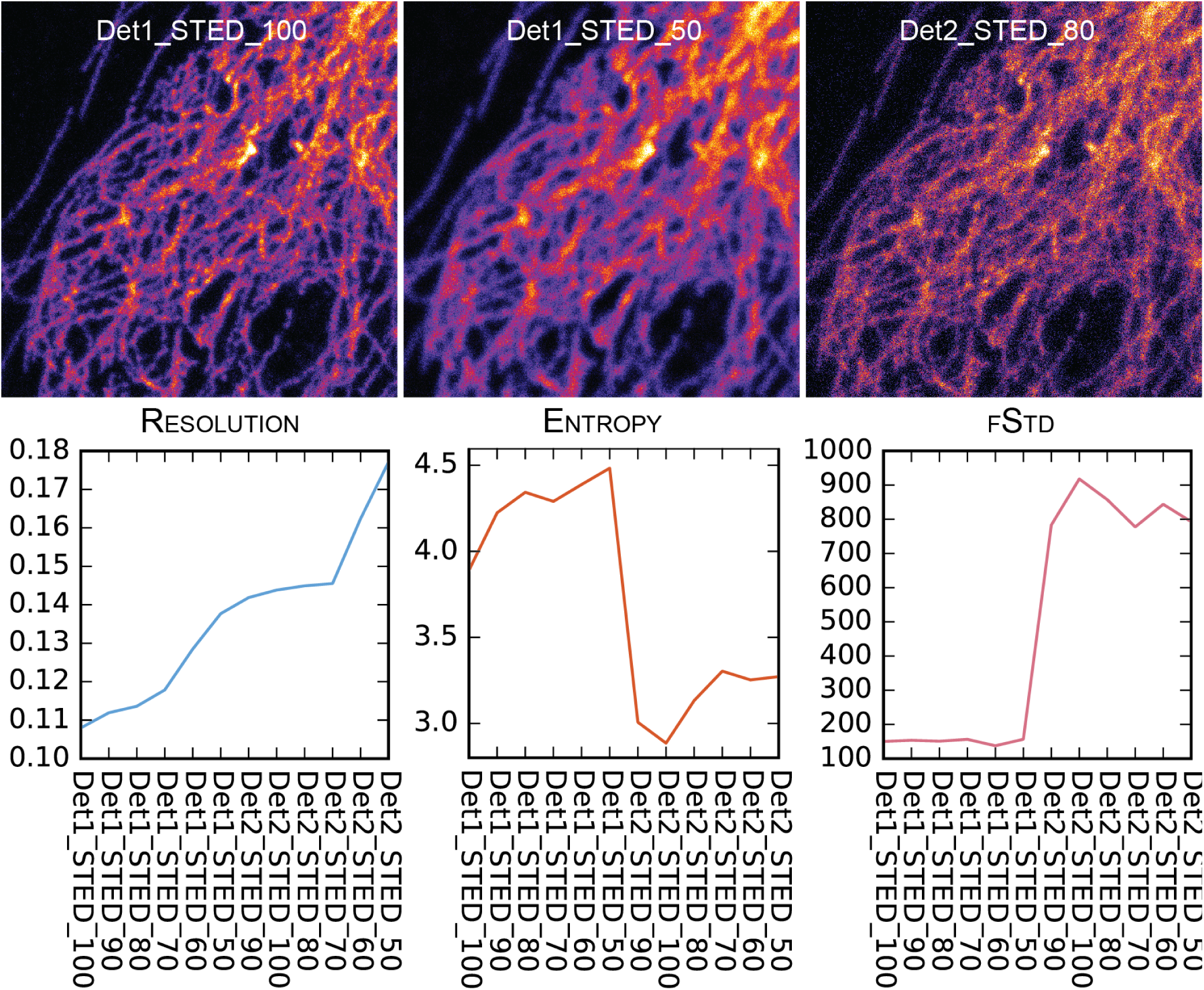
FRC can help in quantitative image quality assessment. The FRC measurement is compared against two image quality parameters on a dataset containing STED super-resolution images with two different detector configurations and five different depletion beam intensities. The spatial domain Entropy measure is very sensitive to contrast changes. The FRC measure gives the effective resolution value. The fSTD measure reacts strongly to noise and blur. The information given by the three measures is very complementary; one might e.g. select a “good” image in the series, based on some sort of combination of these three parameters. [2]

**Table S. 1.** The main advantages and disadvantages of the various splitting methods in terms of sensitivity to spatial bias and temporal correlations are summarized. *yes* is the desired answer here.

| Method | Symmetrical Distribution of Details | Insesitive to fast-axis fluctuations |
| --- | --- | --- |
| a) | yes | yes |
| b) | no | no |
| c) | no | yes |
| d) | yes | no/no |
| e) | no | yes |
| f) | no | no |

## Note S. 1. Compensating for a known image shift in one-image FRC/FSC

The diagonal single image splitting method (Figure S. 4 a)) produces a single pixel shift both in x and y directions. Here we try to understand what such shift does to the FRC/FSC curve. According to the Fourier shift theorem, a shift of (*y*_0_*, x*_0_) pixels of image *f_i_* in the spatial domain, will result into *e*^−*i*2π*sr*/*N*^ frequency dependent phase modulation of its Fourier transform. *r* is equivalent to the radius in the FRC/FSC equation; it should actually be expressed as a vector, but here we abuse the notation a little bit as the summation over rings/spheres in FRC/FSC calculation essentially reduces the dimensionality of a signal to 1D. 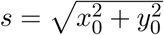 equivalently is the total length of the shift. The FRC/FSC equation can be modified to include the shift as follows:

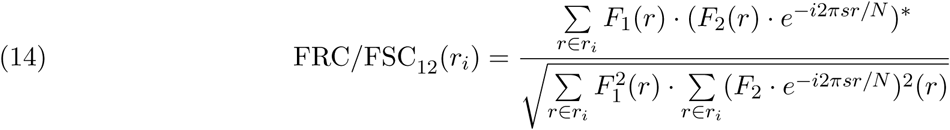

Because (*e*^−*i*2π*sr/N*^)^2^ = 1 it can be altogether ignored in the denominator. On the other hand, because both *F*_1_ and *F*_2_ are Friedel symmetric, the result of summation over Fourier ring/sphere will always be real. The nominator can thus be rewritten as

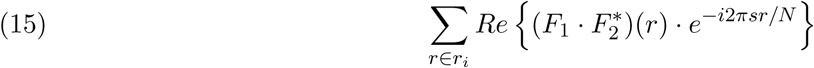

Let’s now make two variable changes 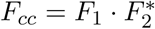 and *D* = *N/*2; substitution to *D* is motivated by the fact that 0 *≤ r_i_ < N/*2. The nominator can now written to form

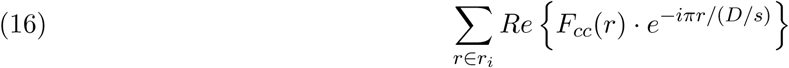

, which ends up being a discrete cosine transform (DCT-I), of length *D/s*. It is known that DCT-I of length *D* is exactly equivalent to a DFT of length 2*D −* 2 with even symmetry. The shift *s* adds a modulation to the DCT that essentially makes the frequency step size larger, thus compressing the spectrum. We compensate for this effect by scaling the pixel size by the shift *s* = 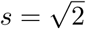.

## Note S. 2. Image descriptions

**Figures 2, 3, 9, S. 5c), S. 1, S. 10, S. 14:** A fixed HeLa cell with alpha-tubulin stained, using Star488 secondary antibodies (Abberior, Germany). Sample was imaged with a Nikon A1 confocal microscope, with excitation wavelength 488nm, 60x/1.4 oil immersion objective and GaAsP detector.The image size is 512×512 and pixel size 51 nm.

**Figures S. 5c), S. 3:** A sample of 40nm Crimson fluorescent nanoparticles (Abberior Instruments, Germany). The sample was imaged with Abberior Instruments Expert Line STED system, excitation wavelength 633nm, depletion wavelength 775nm, 100x/1.4 (UPLSAPO100XO) objective, pinhole 1 Airy unit (AU). Two datasets with two image pairs each (STED, Gated-STED) were acquired with this sample: 800×800, 10nm pixel size; 1000×1000, 15nm pixel size)

**Figure 4a), S. 13, S. 15:** A sample with tubulin cytoskeleton stained with Star-635P (Abberior, Germany). The sample was imaged with Abberior Instruments Expert Line STED system, excitation wavelength 633nm, depletion wavelength 775nm, 100x/1.4 (UPLSAPO100XO) objective. Dimensions (1024×972×30; 30×30×100 nm).

**Figure 4b):** A sample of pollen, imaged with a Nikon A1 confocal microscope, with excitation wavelength 488nm, 40x/1.2 water immersion objective and GaAsP detector. Due to custom intermediate optics on the illumination/detection path, the objective was underfilled, to produce a effective NA of approximately 0.5-0.6.

**Figure S. 16:** A fixed HeLa cell with tubuline cytoskeleton stained with Star635. The sample was imaged with Leica TCS SP5 STED, 100x/1.4 oil immersion objective (HCX PL APO CS), pinhole 1.0AU, excitation wavelength 635nm. The depletion laser wavelength was set to 765nm and its intensity was varied between 50-100% – the same field of view was imaged sequentially with two different detectors. Dimensions (700×700; 15×15nm)

**Figure 1b,d), S. 2, S. 6, S. 7:** A fixed cell with intermediate filaments (vimentin) cytoskeleton stained with Star635. The sample was imaged with Abberior Instruments Expert Line STED system, excitation wavelength 633nm, 100x/1.4 (UPLSAPO100XO) objective. The dataset used in Figures 1b), S. 6 consists of images with different pixel sizes (29 *→* 113 nm; dims 1389×1389 *→* 354×354). The excitation gradient images used in Figure 1d) were imaged with 56nm pixel size (708×708px); the widefield image in Figure S. 7 was acquired with the same parameters, the optical pinhole was opened to remove the optical sectioning effect.

**Figure 1c):** A fixed cell with nuclear pore staining with Star635. The sample was imaged with Abberior Instruments Expert Line STED system, excitation wavelength 633nm, 755nm depletion, 100x/1.4 (UPLSAPO100XO) objective. The STED depletion was raised in 10% steps 0 *→* 20% (arbitrary units) to produce the resolution gradient. Image size (1000×1000; 20×20nm)

**Figure S. 11:** A Atto647N fluorescent layer imaged with bberior Instruments Expert Line STED system, excitation wavelength 633nm, 100x/1.4 (UPLSAPO100XO) objective. Dimensions (500×500; 60×60nm)

## Note S. 3. MIPLIB open-source image analysis software

MIPLIB is an evolution from the *SuperTomo* software [3] that was originally written for the specific purpose of performing multi-view tomographic reconstructions with large 3D STED microscopy datasets. Over the years its scope and features have slowly expanded towards a general purpose microscopy image analysis and processing library. In addition to the new FRC/FSC features, we recently decided to consolidate several previously separate image analysis and processing packages, dealing with e.g. quantitative image quality analysis [2], correlative microscopy [4] and image deconvolution into MIPLIB as well.

The library is being made available under FreeBSD open source license, and it can be downloaded at: https://github.com/sakoho81/miplib

### Regarding iteration in polar/spherical coordinate space

In order to conveniently calculate the (S)FSC measures, flexible methods to index the 3D Fourier space needed to be created. To this end we created a series of spherical-coordinate-system-based Fourier Shell *iterators* in the MIPLIB software. The iterators produce the sequence of 3D indexing structures that are needed to extract voxels on a given shell/section from the Fourier space image for FSC calculation. The iterators were designed to be interchangeable – any one of them can be used in the FSC implementation to produce the desired behaviour. In addition to the regular FSC and SFSC iterators, we also implemented special ones to exclude pixels in certain orientations – this is necessary as microscope images often contain artefacts generated e.g. by the mechanical movement of the xyz - scanning apparatus; the artefact become visible in the FRC/FSC measures as higher than normal resolution values. Excluding pixels in the direction of the optical axis seems to be especially important, as the axial scanning steps (piezo) as well as possible interpolation artefacts otherwise compromise the resolution measure. With each iterator the width of the sections, the thickness of the shells (bin size), as well as the width of the possible exclusion area can be freely selected.

Similar interchangeable iterator scheme is used in the FRC implementation as well, the main difference is that the iterators work in polar coordinate system, and only one cross-correlation curve is calculated for every image. It may in some instances be of interest to exclude certain parts of the Fourier rings as well, if a particular orientation of features is of interest, or if there are artefacts in a certain direction affecting the FRC results.

### Regarding FRC/FSC analysis

The FRC/FSC analysis is a multi-stage process, in which first the image is analyzed to form the FRC/FSC datasets, after which the datasets are analyzed to find out the resolution values. The datasets consist of the cross-correlation histogram and a list of the number of pixels/voxels on each ring/shell; the latter information is needed to calculate the threshold curves. The FRC/FSC analysis workflow is described algorithmically in Algorithm 1.

#### Algorithm 1 Simple pseudocode for our FRC/FSC analysis algorithm. Words in italics denote variable names.

**for** each *Orientation* in *Iterator.Orientations* **do**

calculate FRC/FSC *Dataset*

save to *DataCollection*

**end for**

**for** each *Dataset* in *DataCollection* **do**

fit a curve to *Dataset.Correlation*

calculate *Threshold* from *Dataset.NPoints*

fit curve to *Threshold*

find an intersection of the two curves

**end for**

The resolution thresholds in this paper are based on the (Eq. 3). MIPLIB allows calculating a threshold for any arbitrary SNR_*E*_values, although 0.25 (FRC) and 0.5 (SFSC) are used thoughout this paper. In addition, it is possible to use fixed thresholds (1/7 or any other lever) or the *nσ* thresholds.

The curve fitting has been implemented in two different ways: (I) a standard linear model fitting, with an exponential function of the order of *n* and (II) a piece-wise (smoothed) splines based fitting method. The former method makes it possible to enforce a given shape for the resolution curve, whereas the latter is more robust, especially with very noisy data, into which it is hard to reliably fit a polynomial without some sort of pre-filtering or cropping. We almost exclusively use the smoothed splines as they produce nice looking FRC/FSC curves almost with any kind of data.

The intersection of the cross-correlation and the threshold curves is calculated by minimizing |FRC*/*FSC(*x*) *−* Th(*x*)|, where FRC/FSC(x) and Th(x) are the values of the correlation curve and the threshold curve at frequency *x*, respectively. The optimization is done with a classical Simplex algorithm [5]. In order to reduce the number of iterations, the optimization is started at a frequency *x* at which the FRC/FSC correlation value in the original histogram is just above the threshold curve.

## Note S. 4. Summarizing properties of the single image splitting schemes

Properties of the image splitting methods are summarised. Letters a)-f) refer to the labels used in (Figure S. 4). The main advantages and disadvantages of each method are in addition summarised in (Table S. 1).

a): Diagonal Splitting
  - Since the corresponding pixels of the two images belong to two different line of the original image, artefacts due to temporal correlation are suppressed, i.e the line-dwell time is much longer than the temporal scale of the dynamics.
  - The two generated images are shifted with respect to each other, by the same amount in x and y directions.
b): Horizontal Splitting
  - Since the corresponding pixels of the two images are temporally consecutive artefacts due to temporal correlation maybe introduced.
  - The two generated images are shifted with respect to each other only in the x direction.
  - The x direction is effectively more densely sampled than the y direction
c): Vertical Splitting
  - Since the corresponding pixels of the two images belong to two different lines of the original image, artefacts due to temporal correlation are suppressed; i.e the line-dwell time is much longer than the temporal scale of the dynamics.
  - The two generated images are shifted with respect to each other only in the y direction.
  - The y direction is effectively more densely sampled than the x direction
d): Diagonal Summing
  - Since the corresponding pixels of the two images are obtained adding pixel which belong to the same line of the original image, artefacts due to temporal correlation maybe introduced.
  - The diagonal summing may produce spatial correlations
  - The two images are not shifted with respect to each other.
e): Horizontal Summing
  - Since the corresponding pixels of the two images are obtained by summing pair of temporally consecutive pixel, artefacts due to temporal correlation maybe introduced.
  - The two generated images are shifted with respect to each other only in the y direction.
f): Vertical Summing
  - Since the corresponding pixels of the two images are obtained by summing pixel which belong to two different lines of the original image, artefacts due to temporal correlation are not introduced by the summing; however they may still be present in the split images similar to **b)**.
  - The two generated images are shifted with respect to each other only in the x direction.

